# Cadherins orchestrate specific patterns of perisomatic inhibition onto distinct pyramidal cell populations

**DOI:** 10.1101/2023.09.28.559922

**Authors:** Julie Jézéquel, Giuseppe Condomitti, Tim Kroon, Fursham Hamid, Stella Sanalidou, Teresa Garces, Patricia Maeso, Maddalena Balia, Beatriz Rico

## Abstract

In the mammalian neocortex, excitatory pyramidal cells are assembled into distinct subnetworks, which project to different brain areas. GABAergic interneurons were long thought to connect promiscuously and extensively to pyramidal neurons, but recent evidence supports the existence of a cell type-specific inhibitory connectome. How and when interneurons establish such a precise connectivity pattern among intermingled populations of excitatory neurons remains enigmatic. Here, we investigated the molecular rules shaping cell type- and input-specific inhibitory connectivity in different Layer 5 (L5) pyramidal cell populations. We found that neighboring L5 intra- (L5 IT) and extra-telencephalic (L5 ET) neurons receive different combinations of inhibitory perisomatic inputs from Parvalbumin- (PV+) and Cholecystokynine-positive (CCK+) basket cells. We also identified *Cdh12* and *Cdh13*, two cadherin superfamily members, as critical mediators of L5 pyramidal cell type-specific inhibitory connectivity. Our data revealed a minimal overlap between L5 IT and L5 ET presynaptic inhibitory networks, and suggests that different populations PV+ basket cells innervate distinct L5 pyramidal cell types. Altogether, our work unravels the contribution of cadherins in shaping cortical interneuron wiring and provides new insights into the development of inhibitory microcircuits.

## Introduction

Precise and fine-tuned neuronal connectivity underlies even the most basic animal behavior. This connectivity reaches an extreme complexity in the mammalian cerebral cortex, where two main types of neurons coexist: glutamatergic principal or pyramidal cells and gamma-aminobutyric acid-expressing (GABAergic) interneurons. Pyramidal cells are highly heterogeneous and are organized into distinct neuronal ensembles, forming unique information-processing streams ^1,2^. By selectively targeting different subcellular compartments of principal cells, inhibitory interneurons play a critical role in gating cortical activity and ensuring proper brain computation ^3^. However, how interneurons regulate such intricate and diverse excitatory networks remains enigmatic.

The classical vision assumed that interneurons indiscriminately target all pyramidal cells in the cerebral cortex ^4–6^. However, there is now substantial evidence indicating that cell type-specific connectivity motifs mediate the wiring of inhibitory connections ^7–12^. These studies suggest instead that interneurons discriminate among different postsynaptic targets and exhibit biased connectivity patterns ^7,8,10,12–15^. For example, pyramidal cells projecting to subcortical areas receive more perisomatic inhibition from parvalbumin-expressing (PV+) basket cells than local-projecting neighboring cells ^8,14^. Similarly, cholecystokinin-expressing (CCK+) basket cells preferentially target pyramidal cells sending axons outside the hippocampus ^7^. Consequently, interneurons not only influence pyramidal cell activity by targeting different subcellular compartments but can also gate the information flow of distinct subnetworks of pyramidal cells.

How interneurons select specific targets during development and what molecular programs control such refined connectivity motifs to arise is unknown. Here, we took advantage of the stereotypical organization of cortical layer 5 (L5) and the coexistence of two pyramidal cell types with distinct projection targets to explore the emergence and molecular determinants of cell type-specific perisomatic inhibition. Focusing on L5 intra- (L5 IT) and extra-telencephalic (L5 ET) pyramidal cells ^16–20^, we demonstrate that these pyramidal cell populations receive different ratios of PV+ and CCK+ inputs. We also found that members of the Cadherin superfamily, *Cdh12* and *Cdh13*, contribute to instructing cell type- and input-specific perisomatic inhibition. Moreover, simultaneous mapping of the inhibitory inputs targeting L5 IT and L5 ET revealed the contribution of different populations of PV+ basket cells to individual pyramidal cells. Altogether, our findings unveil a molecular and cellular code orchestrating the assembly of inhibitory microcircuits targeting different pyramidal cell populations.

## Results

### Differential perisomatic inhibition onto distinct pyramidal cell types

Previous studies in the hippocampus, entorhinal cortex and prefrontal cortex have shown that the levels of perisomatic inhibition differ between pyramidal cells with distinct projection targets ^7–11^. To investigate whether this principle is conserved in other cortical areas, we took advantage of the stereotypical organization of layer L5 in the primary somatosensory cortex (S1), where distinct pyramidal cell types can be segregated based on their axonal projections ^16^. In this region, L5 IT neurons send their axons to the contralateral hemisphere and other cortical areas, while L5 ET neurons target subcortical structures, including the thalamus, superior colliculus, and brainstem nuclei ^19^. To simultaneously label L5 IT and L5 ET neurons, we co-injected RB488 and RB555 fluorophore-coated latex retrobeads in the contralateral S1 and the ipsilateral pons of wild-type neonates (Fig. 1a and Extended Data Fig. 1). We chose cortico-pontine projecting neurons as a reference for L5 ET neurons because early postnatal injections in the pons provided the most reliable labeling among the different subcortical areas tested. Retrograde tracing from the injection sites revealed a stereotypical double-layer cell distribution (Fig. 1a) whereby L5 ET neurons were restricted to L5b while L5 IT cells were enriched in L5a but present across the entire layer (Fig. 1a). To determine whether these two populations of neurons receive different levels of perisomatic inhibition, we recorded miniature inhibitory postsynaptic currents (mIPSCs) from neighboring retrogradely labeled S1 L5 IT and L5 ET neurons. Whole-cell voltage-clamp recordings at P30 revealed that the mIPSC frequency was higher in L5 ET than in L5 IT neurons, whereas both cell types exhibited similar mIPSC amplitude (Fig. 1b). These results reveal that S1 L5 ET cells receive significantly more inhibitory inputs than L5 IT neurons.

**Fig. 1.**
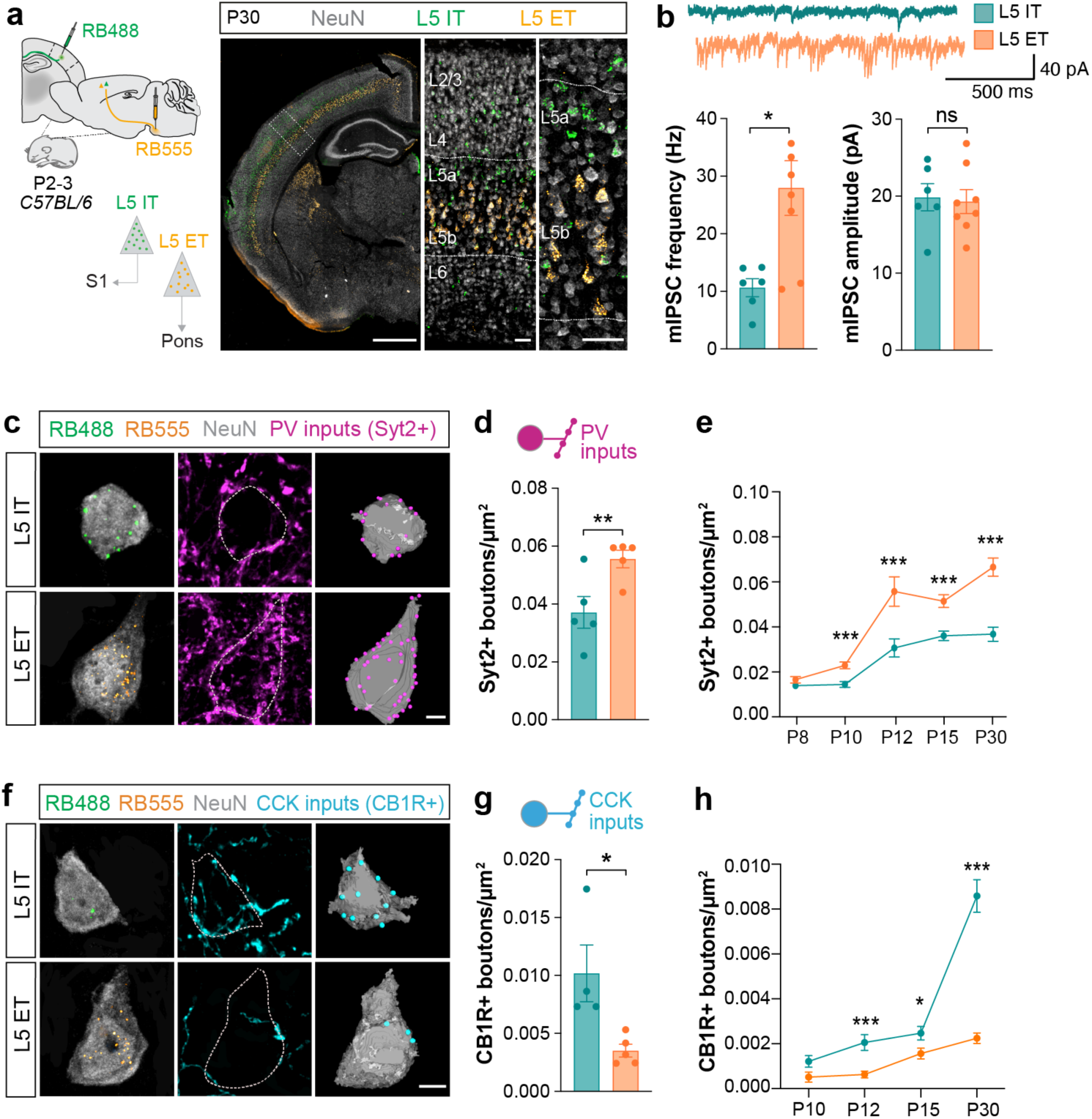
Differential perisomatic inhibition onto distinct pyramidal cell types. **a,** L5 IT and L5 ET neurons retrograde labelling strategy. Green (RB488) and red (RB555, pseudocolor in orange) fluorescent retrobeads were injected in the contralateral somatosensory cortex (S1) and in the ipsilateral pons of P2-3 C57BL/6J mice to target L5 IT and L5 ET axons, respectively. Pyramidal cell somas were labeled with NeuN. L5 IT neurons (green) were mainly enriched in L5a, while L5 ET (orange) were exclusively located in L5b. **b,** mIPSCs frequency (Hz) (*L5 IT* n = 6 cells from 3 mice, *L5 ET* n = 8 cells from 2 mice; Mann-Whitney test * *P* < 0,05) and mIPSCs amplitude (pA) (*L5 IT* n = 6 cells from 3 mice, *L5 ET*; n = 8 cells from 2 mice; ns, Mann-Whitney test) from retrogradely-labeled L5 IT and L5 ET cells at P30. **c,** Confocal and 3D-reconstructed images illustrating PV inputs (Syt2+, magenta) onto L5 IT and L5 ET somas labelled with NeuN (grey) and retrobeads. **d,** Syt2+ boutons density onto L5 IT and L5 ET at P30 (*L5 IT* n = 5 mice, 62 cells, *L5 ET* n = 5 mice, 92 cells; paired t-test ** *P* < 0.01). **e,** Syt2+ boutons density onto L5 IT and L5 ET over post-natal development (*P8 L5 IT* = 60 cells, *L5 ET* = 116 from 5 mice; *P10 L5 IT* = 131 cells, *L5 ET* = 123 cells from 6 mice; *P12 L5 IT* = 103 cells, *L5 ET* = 109 cells from 5 mice; *P15 L5 IT* = 164 cells, *L5 ET* = 125 cells, 7 mice; *P30 L5 IT* = 61 cells, *L5 ET* = 97 cells from 5 mice; multiple t-test Holm-Šídák posthoc, P8 ns, P10 \*\*\**P* < 0.001, P12 \*\**P* < 0.01, P15 \*\*\**P* < 0.001, P30 *** *P* < 0.001). **f,** Confocal and 3D-reconstructed images illustrating CCK inputs (CB1R+, cyan) onto L5 IT and L5 ET somas labelled with NeuN (grey) and retrobeads at P30. **g,** CB1R+ boutons density onto L5 IT and L5 ET at P30 (*L5 IT* n = 4 mice, 82 cells, *L5 ET* n= 5 mice, 83 cells; t-test \**P* < 0.05). **h,** CB1R+ boutons density onto L5 IT and L5 ET over postnatal development (*P10 L5 IT* n = 73 cells, *L5 ET* n = 45 cells from 5 mice; *P12 L5 IT* n = 57 cells, *L5 ET* n = 49 cells from 3 mice; *P15 L5 IT* n= 107 cells, *L5 ET* n = 80 cells, from 5 mice; *P30 L5 IT* n = 127 cells, *L5 ET* n = 120 cells from 5 mice; multiple t-test Holm-Šídák posthoc: P10 ns, P12 \*\*\**P* < 0.001, P15 \**P* < 0.05, P30 \*\*\**P* <0.001). Data are represented as mean ± s.e.m, \**P* < 0.05; \*\**P* < 0.01; \*\*\**P* < 0.001; ns, not significant. Each dot represents an individual mouse. Data are mean ± s.e.m. Scale bars, 500 μm, 50 μm (a) and 5 μm (b, c).

### Cell type-specific inhibitory motifs onto pyramidal cell populations

PV+ and CCK+ basket cells are the two major sources of perisomatic inhibition in the cerebral cortex ^21,22^. To investigate the contribution of each interneuron population to the perisomatic inhibition of L5 pyramidal neurons, we quantified the density of PV+ and CCK+ synapses onto the soma of L5 IT and L5 ET neurons at P30 using specific presynaptic markers: Synaptotagmin 2-positive (Syt2+) for PV+ inputs ^23^ and cannabinoid receptor 1 (CBR1) for CCK+ inputs ^24^. We found that PV+ boutons were twice more abundant in L5 ET neurons than in L5 IT cells (Fig. 1c, d), confirming our functional analysis (Fig. 1b). In contrast, L5 IT neurons received more CCK+ boutons labelled with CB1R than L5 ET cells (Fig. 1c and 1g). Given that pyramidal cells receive considerably more PV+ than CCK+ inputs (^25^ and present study), our mIPSC recordings might mainly reflect PV+ innervation, masking the contribution of CCK+ basket cells (Fig. 1b).

We next investigated when these distinct patterns of perisomatic inhibition emerge. To this end, we characterized the developmental timeline of PV+ and CCK+ inputs onto L5 IT and L5 ET neurons. We observed that the differences in PV+ innervation already existed during early stages of synapse formation (P10) and increased throughout postnatal development (Fig. 1e). CCK+ inhibition followed a slower progression, although the preferential targeting of L5 IT neurons was already evident by P12 and got gradually more robust over time (Fig. 1h). Altogether, our results indicate that PV+ and CCK+ interneurons provide cell type-specific innervation onto neighboring L5 pyramidal cell populations embedded in distinct neuronal networks, and that such connectivity patterns are determined during the early stages of synapse formation.

### Cell type-specific transcriptional programs in distinct L5 pyramidal cells

What molecular mechanisms underlie the different innervation patterns of PV+ and CCK+ interneurons? We hypothesized that the postsynaptic targets (i.e. L5 IT and L5 ET neurons) can shape interneuron inputs to match their inhibitory requirements by expressing cell type-specific synaptic programs. We examined the transcriptomic signatures of L5 IT and L5 ET neurons when the differential perisomatic inhibition patterns first emerged to explore whether postsynaptic molecules could instruct cell type-specific levels of perisomatic inhibition. We dissected S1 deep layers at P10 and isolated retrogradely labeled L5 IT and L5 ET neurons using fluorescence-activated cell sorting. We then performed bulk RNA-sequencing (RNAseq) and compared the gene expression profiles of both populations of pyramidal cells (Fig. 2a and Extended Data Fig. 2a). Using this approach, we identified multiple cell type-specific genes, including known markers of L5 IT neurons such as *Lmo4* and *Satb2*, and of L5 ET cells like *Bcl11b* and *Crym* (Extended Data Fig. 2b). Although astroglia or oligodendroglia genes were absent from our dataset, we unexpectedly found some microglial genes in the L5 ET population (Extended Data Fig. 2c). To enrich our study with L5-specific genes, we used bioinformatics to intersect our dataset with two other RNA-seq datasets ^26,27^ (Fig. 2b and Extended Data Fig. 2d). In brief, genes differentially expressed (DEGs) in our dataset and in ^26^ were filtered for neonatal-regulated genes ^27^. A final list of DEGs was then identified by combining: (1) the two studies (130 DEG), (2) the present study only (342 DEG), and (3) Klingler et al., 2019 only (381 DEG) (Fig. 2c). Following this criterion, we obtained 853 DEG between L5 IT and L5 ET neurons (Fig. 2c-e, and Extended Data Fig. 2e, f). Gene Ontology (GO) analysis of this restricted dataset unveiled enrichment in genes involved in the organization and function of the synapse (Fig. 2e). L5 IT-enriched genes contributed to a higher proportion of these synaptic GO terms, but terms relevant to perisomatic inhibition, such as “GABAergic synapse” and “cell-cell adhesion” also featured in the analysis were equally represented in L5 IT and L5 ET cells (Extended Data Fig. 2e).

**Fig. 2.**
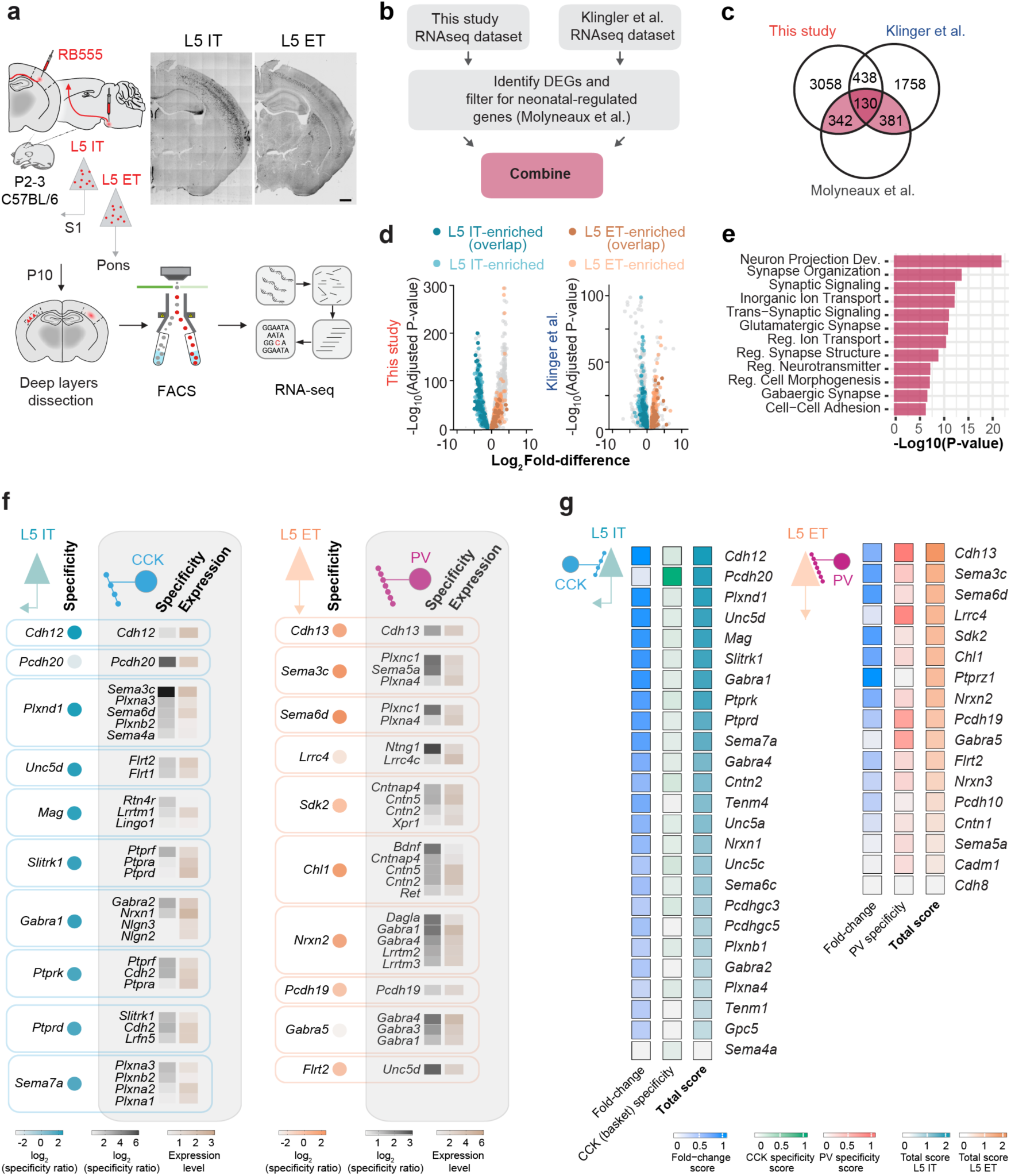
Cell type-specific transcriptional programs in L5 pyramidal cells subpopulations. **a**, Experimental design. **b,** Intersection strategy of our RNAseq dataset with two other studies. **c,** Differentially expressed genes (DEGs) between L5 IT and L5 ET neurons in the three RNAseq datasets. **d,** Volcano plot of DEGs in our study and Klingler et al. **e,** Gene ontology terms significantly enriched in the final dataset. **f.** Cell-specific candidate cell-surface molecules expressed in L5 IT and L5 ET neurons ranked based on their putative pre-synaptic partners’ specificity and expression level in CCK basket cells and PV interneurons, respectively. This analysis relies on known protein-protein interactions and gene expression in CCK+ and PV+ cell types described as such, according to (Yao et al., 2021). **g,** Heatmaps showing the fold-change expression, specificity score and resulting total score for L5 IT and L5 ET DEG coding for cell-surface molecules. Scale bar, 500 μm (a).

Several families of cell-surface molecules have been implicated in cell-cell type recognition during neural circuit development ^28–31^. We then curated the DEGs list for genes coding cell-surface molecules to identify candidate genes instructing cell type-specific perisomatic inhibition in L5 pyramidal cell populations. Genes were ranked according to the sum of a specificity score measuring the expression of a corresponding cell-surface partner in the preferential presynaptic targeting interneuron (i.e., CCK-L5 IT and PV-L5 ET, Fig. 2f) - and the gene expression fold-change score (Fig. 2g). Using this set of criteria, two cadherin superfamily members appeared as top candidate genes – Cadherin-12 (*Cdh12*) for L5 IT and Cadherin-13 (*Cdh13*) for L5 ET. To validate our bioinformatic predictions, we performed single-molecule RNA fluorescent in situ hybridization and measured *Cdh12* and *Cdh13* transcript levels in both populations of L5 pyramidal neurons at P10 (Extended Data Fig. 3). We retrogradely labeled L5 IT and L5 ET neurons in these experiments using adeno-associated viruses (AAVs) expressing tag-BFP or tdTomato, respectively (Extended Data Fig. 3a). *Cdh12* RNA expression was three times more enriched in L5 IT than L5 ET neurons (Extended Data Fig. 3b, c). Conversely, *Cdh13* RNA was twice more abundant in L5 ET neurons than in L5 IT neurons (Extended Data Fig. 3d, e). These findings reveal that L5 IT and L5 ET express a unique cadherin signature, which mirrors cell type-specific PV+ and CCK+ basket cell connectivity motifs onto L5 pyramidal cell populations.

### Cell type- and input type-specific inhibition of pyramidal cell populations require *Cdh12* and *Cdh13*

To assess whether postsynaptic *Cdh12* and *Cdh13* play a role in instructing CCK+ and PV+ basket cell inputs in the two populations of pyramidal cells, we performed cell type-specific loss-of-function experiments using a conditional knockdown (KD) strategy *in vivo* ^32^. We designed Cre-dependent AAVs expressing short-hairpin RNAs against control (*shLacZ*) and our candidate genes (*shCdh12* and *shCdh13*) and confirmed their efficiency *in vitro* (Extended Data Fig. 4a, b). To specifically down-regulate the expression of our candidate genes in L5 IT neurons (Fig. 3a), we used the *Tlx3-Cre* mouse line, which labels these cells^33^. To target L5 ET cells, we injected a retrograde Cre-expressing AAV virus (*rAAV2retro-Cre*) in the pons of wild-type mice. In both cases, we injected Cre-dependent *shCdh12* and *shCdh13* AAV in S1 (Fig. 3e). We confirmed the efficiency of both approaches to down-regulate the expression of *Cdh12* and *Cdh13 in vivo* (Extended Data Fig. 4c-f) and quantified the density of PV+ and CCK+ inputs on the somas of L5 IT and L5 ET neurons at P30. We found that *Cdh12* KD did not impact the PV+ innervation of L5 IT neurons (Fig. 3b, c). In contrast, *Cdh12* KD significantly reduced CCK+ inputs onto L5 IT cells, which remained unaffected following *Cdh13* KD (Fig. 3b, d). Conversely, *Cdh13* KD reduced the number of PV+ inputs targeting L5 ET neurons without affecting the CCK+ inhibition (Fig. 3f-h). Consistent with these observations, we found that the *Cdh13* KD reduced mIPSCs frequency in L5 ET neurons (Extended Data Fig. 5). *Cdh12* KD did not reveal changes in mIPSCs frequencies in L5 IT neurons (Extended Data Fig. 5), which may be explained by the small contribution of CCK+ inputs compared to PV+ inputs to the total inhibition received by these cells. Surprisingly, *Cdh12* KD did not impact CCK+ inputs targeting L5 ET neurons, and *Cdh13* KD did not modify PV+ inhibition in L5 IT pyramidal cells (Fig. 3b, c, f, h), which suggests a cell type-specific role for *Cdh12* and *Cdh13* in the wiring of CCK+ and PV+ inputs. Interestingly, our results also revealed that reducing PV+ or CCK+ inputs in L5 pyramidal cells does not trigger an apparent compensation from the other inhibitory source. This observation is consistent with the idea that these two types of interneurons play different roles in modulating the activity of pyramidal cells. Altogether, our findings suggest that *Cdh12* expression in L5 IT neurons instructs CCK+ innervation, whereas *Cdh13* expression in ET neurons recruits PV+ inputs.

**Fig. 3.**
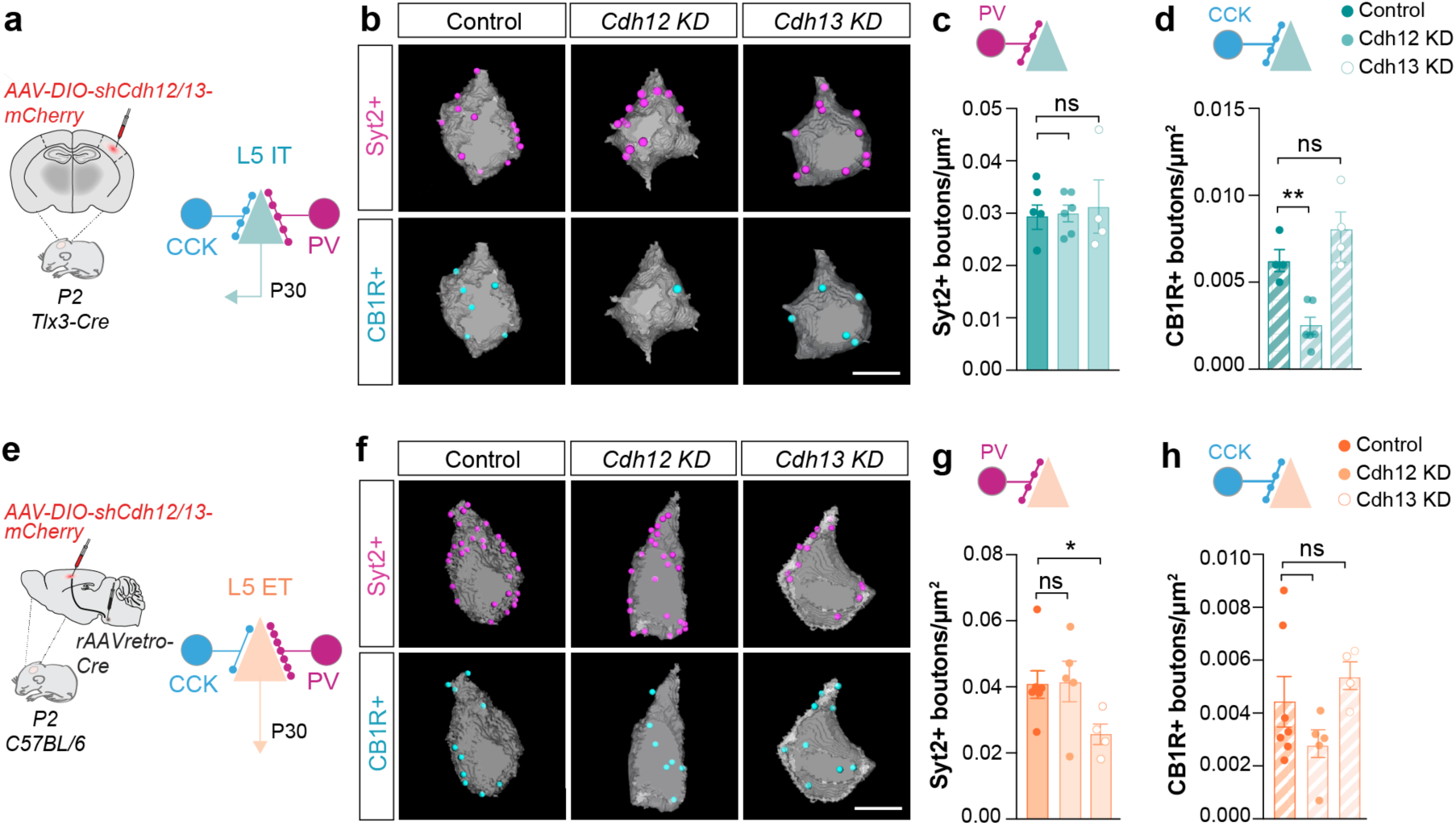
Cadherins drive cell type- and input-specific inhibition onto L5 pyramidal cell subpopulations. **a,** Conditional viral strategy to down-regulate *Cdh12* and *Cdh13* in L5 IT cells and measure its impact on perisomatic inputs at P30. **b,** 3D-reconstructed confocal images of PV+ (Syt2+, magenta) and CCK+ inputs (CB1R+, cyan) onto L5 IT somas. **c,** Syt2+ boutons density onto L5 IT (*Control* n = 4 mice, 94 cells; *Cdh12 KD* n = 6 mice, 140 cells; *Cdh13 KD* n = 4 mice, 119 cells; One-way ANOVA ns, Tukey’s posthoc: Control vs *Cdh12* KD ns, Control vs *Cdh13* KD ns, *Cdh12* KD vs *Cdh13* KD ns). **d,** CB1R+ boutons density onto L5 IT (*Control* n = 4 mice, 95 cells; *Cdh12 KD* n = 6 mice,141 cells; *Cdh13 KD* n = 4 mice, 107 cells; One-way ANOVA \*\*\**P* < 0.001, Tukey’s posthoc: Control vs *Cdh12* KD \*\**P* < 0.01, Control vs *Cdh13* KD ns, *Cdh12* KD vs *Cdh13* KD \*\*\**P* < 0.001). **e,** Conditional viral strategy to down-regulate *Cdh12* and *Cdh13* in L5 ET and the subsequent quantification of perisomatic inputs at P30. **f,** 3D-reconstructed confocal images of PV+ (Syt2+, magenta) and CCK+ inputs (CB1R+, cyan) onto L5 ET somas. **g,** Syt2+ boutons density onto L5 ET (*Control* n = 9 mice, 232 cells; *Cdh12 KD* n *=* 5 mice, 114 cells; *Cdh13 KD* n = 6 mice, 127 cells; One-way ANOVA *P < 0.05, Tukey’s posthoc: Control vs *Cdh12* KD ns, Control vs *Cdh13* KD \**P* < 0.05, *Cdh12* KD vs *Cdh13* KD \**P* < 0.05). **h,** CB1R+ boutons density onto L5 ET (*Control* n = 9 mice, 163 cells); *Cdh12 KD* n = 4 mice (95 cells); *Cdh13 KD* n = 6 mice, 98 cells; One-way ANOVA ns, Tukey’s posthoc: Control vs *Cdh12* KD ns, Control vs *Cdh13* KD ns, *Cdh12* KD vs *Cdh13* KD ns). \**P* < 0.05; \*\**P* < 0.01; \*\*\**P* < 0.001; ns, not significant. Each dot represents an individual mouse. Data are represented as mean ± s.e.m. Scale bar, 10μm (b, f).

### Mapping pyramidal cell populations inhibitory connectivity

Our previous experiments revealed that *Cdh12* and *Cdh13* down-regulation does not affect perisomatic inputs in the same way in the two populations of L5 pyramidal cells (Fig. 3). This suggests that the same molecule, expressed in different cell types, may not instruct the same connectivity motifs. A plausible hypothesis is that distinct molecular codes exist for each population of pyramidal cells because interneuron populations are also diverse. Consistent with this notion, interneuron diversity is extensive, and several subtypes of PV+ and CCK+ basket cells have been identified based on their morphological, electrophysiological and transcriptomic (MET-types) signatures ^34^. We thus hypothesized that different subtypes of PV+ and CCK+ interneurons innervate L5.

To test this hypothesis, we engineered a multiplex rabies-based monosynaptic tracing approach ^35,36^ to delineate the cell type-specific inhibitory input maps of each population of L5 pyramidal cells (Fig. 4a). In these experiments, we focused on PV+ basket interneurons because they represent the most abundant source of perisomatic inhibition in the cerebral cortex ^37–40^. Multiplex monosynaptic tracing relies on specific rabies viral envelopes (EnvX) and their corresponding receptors (TVX). To simultaneously visualize presynaptic networks of L5 IT and L5 ET neurons, we designed distinct *EnvA/TVA* and *EnvB/TVB* complexes and employed two recombinase systems to selectively target L5 IT and L5 ET cells in the same animal (Fig. 4a). We injected a *FlpO-*expressing retrograde AAV in the pons of *Tlx3-Cre* mice; hence neighboring L5 ET and L5 IT neurons would express *FlpO* and *Cre*, respectively (Fig. 4a-c). We injected Flp- and Cre-dependent viruses in S1 to obtain cell type-specific receptor expression. To target L5 ET neurons, we injected two *FlpO*-dependent AAVs, one encoding a mutant TVA and a membrane-bound GFP and another encoding the RV glycoprotein *oG* (Fig. 4a-d). We then injected *Cre-*dependent AAVs expressing a BFP-tagged TVB and *oG* to target L5 IT cells (Fig. 4a-d). Three weeks post-injections, we injected two distinct rabies into S1, one encoding G-deficient EnvA and tdTomato and another encoding G-deficient EnvB and nuclear GFP (Fig. 4a-g and Extended Data Fig. 6). This combination allowed retrograde monosynaptic tracing and visualization of inputs in both populations of L5 pyramidal cells, which we examined after the rabies injection.

**Fig. 4.**
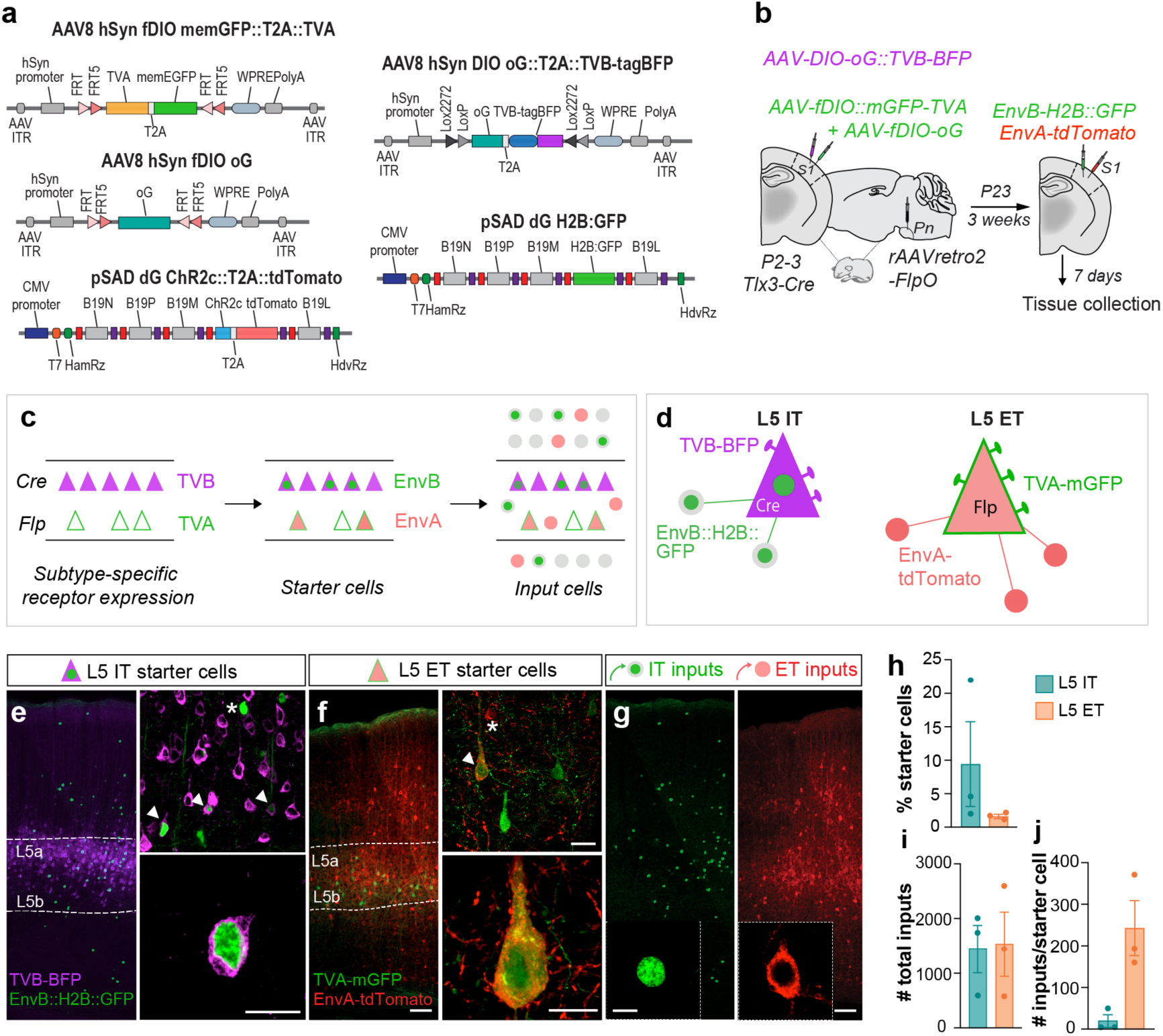
Mapping pyramidal cell subpopulations inhibitory connectivity. **a,** Constructs of the viral *Cre*- and *FlpO*-dependent oEnvA/oTVA and oEnvB/oTVB constructs. **b,** Multiplex monosynaptic tracing strategy. A *FlpO*-expressing retrograde AAV was injected into the pons (Pn) of P2-3 *Tlx3-Cre* mice. A cocktail of *Cre*-dependent oG and TVB-BFP expressing AAV, together with *FlpO*-dependent oG and TVA-mGFP AAVs were injected into the S1. Three weeks later, *EnvA-RVΔG-tdTomato* and *EnvB-RVΔG-H2B:GFP* were co-injected into the S1. **c,** Schematic of the spatial distribution of receptor-expressing cells, starter cells and input cells for L5 IT and L5 ET populations. Note that L5 IT cells present in L5a express the *Cre* recombinase and can be infected with the TVB-receptor (magenta triangle) and/or EnvB (green and magenta triangle). L5 ET cells present in L5b express the *FlpO* recombinase and can be infected with the TVA-receptor (green outlined triangle) and/or EnvA (green and red triangle). Inputs from L5 IT (green circle) and L5 ET (red circle) neurons can then be mapped across the different cortical layers. **d,** Schematic representation of L5 IT and L5 ET starter cells and their direct inputs expressing the different viruses. **e,** Confocal images illustrating the spatial distribution of L5 IT starter cells. Top panel: L5 IT starter cells (white arrowhead) co-expressing TVB-BFP (magenta) and EnvB-H2B:GFP (green) were intermingled in L5 with TVB-BFP+ (asterisk) and EnvB-H2B:GFP-expressing cells. Bottom panel: Confocal image illustrating a L5 IT starter cell. **f,** Confocal images illustrating the spatial distribution of L5 ET starter cells. Top panel: L5 ET starter cells (white arrowhead) co-expressing TVA-mGFP (green) and EnvA-tdTomato (red) could be detected in L5b, among TVA-mGFP expressing cells and EnvA-tdtTomato expressing cells (asterisk). **g,** Confocal images illustrating the distribution of L5 IT (green) and L5 ET (red) input cells across a cortical column. L5 IT inputs were labelled with a nuclear GFP while L5 ET inputs expressed a cytosolic tdTomato. **h,** Fraction of L5 IT and L5 ET starter cells (n = 3 mice). **i,** Number of L5 IT and L5 ET total inputs (n = 3 mice). **j,** Number of L5 IT and L5 ET inputs per starter cell (n = 3 mice). Data are represented as mean ± s.e.m. Scale bars, 50 μm (e, f, g), 25 μm (e, f), 5 μm (g).

We observed that L5 IT starter cells (expressing BFP and nuclear GFP) were mainly located in L5a (Fig. 4c, e), whereas L5 ET starter cells (expressing membrane-bound GFP and cytosolic tdTomato) were restricted to L5b (Fig. 4c, f). It is worth mentioning that the two recombination systems used in this approach are not equally efficient because *Cre* expression was achieved in virtually all L5 IT neurons *via* the *Tlx3-Cr*e mouse line, while variable *Flp* viral copies were expressed in a fraction of L5 ET cells. We consistently observed more L5 IT starter cells than L5 ET starter cells (Fig. 4e, f, h). Nevertheless, the total number of inputs reaching each L5 pyramidal cell type was comparable (Fig. 4h), validating our experimental approach. Furthermore, we did not observe any labeled cells in wild-type mice, further confirming the specificity of our method (Extended Data Fig. 6). We then mapped the afferent connections of L5 IT and L5 ET starter cells throughout the neocortex. In each brain, we quantified the number of inputs for each L5 pyramidal cell type as a fraction of the total number of inputs in the entire neocortex (Fig. 4i). We observed that the number of inputs per starter cell was much higher for L5 ET neurons than L5 IT cells (Fig. 4j), consistent with other studies showing that L5 ET neurons receive more neuronal connections than L5 IT neurons ^41^.

To compare the fraction of inhibitory inputs onto both L5 pyramidal cell populations, we then quantified the proportion of PV+ inputs over the total number of inputs in the neocortex (Fig. 5a-c). We found that the fraction of PV+ input cells targeting L5 ET neurons was significantly higher than in L5 IT cells (Fig. 5b). These results were consistent with our synaptic analysis and confirmed that L5 ET neurons receive more PV+ inputs from more PV+ cells than L5 IT neurons. We also examined whether the same PV+ basket cells could innervate both populations of L5 pyramidal cells or whether different PV+ cells target specific L5 ET and L5 IT neurons. In the first scenario, we would expect a significant fraction of PV+ cells containing both rabies. Otherwise, most PV+ cells should only express nuclear GFP or tdTomato. We found a minimal fraction of double-labeled afferent PV+ cells (Fig. 5a, b, d), demonstrating that most PV+ cells innervated either L5 IT neurons or L5 ET cells. The very few PV+ cells contacting both populations of L5 pyramidal cells were primarily allocated in layer 5a (Fig. 5e). Moreover, we found that the PV+ interneurons contacting L5 IT and L5 ET cells have distinct laminar distributions (Fig. 5e). We observed that PV+ interneurons projecting to L5 IT neurons were spread across L2/3, L4 and L5a, while L5 ET neurons primarily received local inhibition from L5a and L5b PV+ cells (Fig. 5e). In sum, our experiments suggest that L5 IT and L5 ET neurons receive inhibition from segregated presynaptic networks of PV+ cells.

**Fig. 5.**
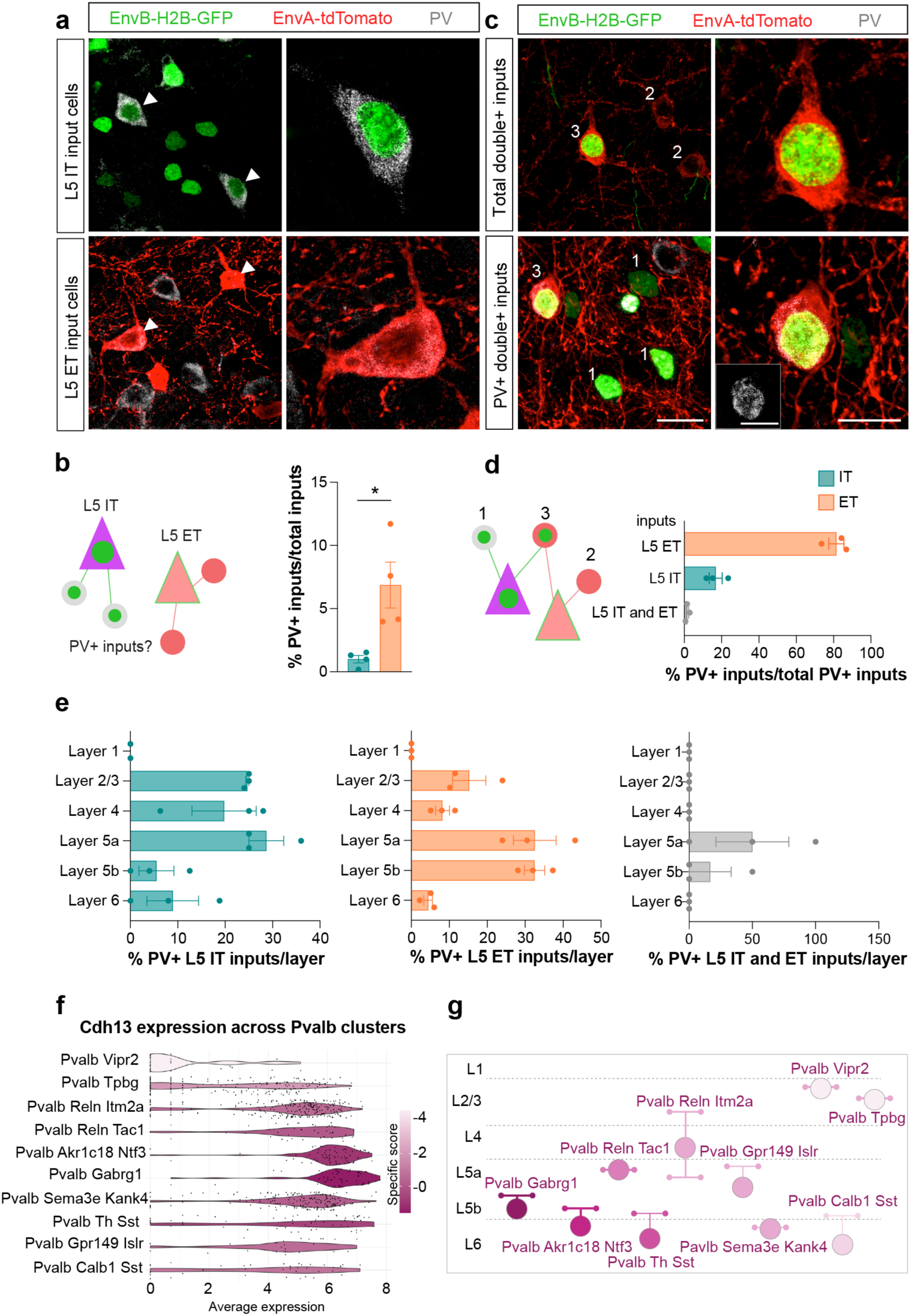
Different PV+ subtypes innervate L5 IT and L5 ET pyramidal cell types. **a,** Confocal images illustrating L5 IT (green) and L5 ET (red) input cells positive for PV staining (grey). PV+ input cells are indicated with a white triangle. **b,** Fraction of PV+ inputs among L5 IT and L5 ET input cells (n = 3 mice). **c,** Confocal images illustrating L5 IT (type 1, EnvB-H2B:GFP) L5 ET (type 2, EnvA-tdTomato) or input cells infected with the two rabies types (type 3, green and red). A minimal fraction of input cells infected with EnvA and EnvB rabies were also positive for PV (grey). **d,** Proportion of PV+ inputs among L5 IT, L5 ET or L5 IT/ET inputs (n = 3 mice). **e,** Proportion of PV+ inputs per layer for L5 IT, L5 ET and L5 IT/ET inputs (n = 3 mice). **f,** Violin plot of *Cdh13* RNA expression and specificity score across the different PV+ MET-types described by the Allen Brain Atlas (Gouwens et al., 2020). **g,** Schematic laminar distribution of the different PV+ MET-types color-coded for *Cdh13* expression levels. Data are represented as mean ± s.e.m. Scale bars, 20μm (a, c), 10μm (a, c).

Based on these findings, we hypothesized that different interneuron types may target specific populations of pyramidal cells by expressing matching molecular programs. To address this question, we analyzed *Cdh13* expression levels across different types of PV+ interneurons ^34^. We found that PV+ cells with their somas or axonal arbor in L5b (*Pvalb-Gabrg1* and *Pvalb-Akr1c18-Ntf3*) contain the highest levels of *Cdh13* mRNA, whereas L5a PV+ interneurons (*Pvalb-Reln-Tac1* and *Pvalb-Gpr149-Islr*) have the lowest levels of *Cdh13* (Fig. 5f, g).

In summary, our findings support the notion that perisomatic inhibition targeting different populations of L5 pyramidal cells follows strictly defined rules. Firstly, there is a clear preference for the source of presynaptic inhibitory cells (i.e., PV+ versus CCK+ cells). Secondly, there seems to be a precise allocation of specific subtypes of interneurons (i.e., distinct PV+ MET-types) for each type of L5 pyramidal cell.

## Discussion

Understanding the core principles governing neuronal connectivity is essential to identifying the intricate flow of information across neuronal circuits in the cerebral cortex. Cortical interneurons exhibit a remarkable diversity and are critical for sculpting the information conveyed by pyramidal cells. Over the last years, new evidence has transformed our view on inhibition, shifting from promiscuous connectivity ^4–6,42^ to a precise pattern of interneuron connections whose rules are only beginning to be understood ^7–12^. However, how this inhibition emerges, and the molecular programs underlying the emergence of precise cell-cell recognition is still unknown. Our study demonstrates that pyramidal cells shape interneuron wiring by expressing specific molecular codes that shape cell type-specific inhibitory connectivity. We found that distinct populations of L5 pyramidal cells promote unique perisomatic inhibitory patterns by expressing two cadherins. *Cdh12* conducts CCK+ inhibition onto L5 IT neurons, and *Cdh13* exclusively controls PV+ inhibition onto L5 ET cells. This differential inhibition pattern is acquired by L5 pyramidal cells during the initial steps of synaptogenesis, suggesting that connectivity features might be intrinsically imprinted during cell type specification.

The amount of inhibition received by pyramidal cells correlates with their projection pattern ^7–12^. Our findings demonstrate that this rule is conserved across different cortices, revealing a consistent preference of L5 IT neurons for CCK+ inputs L5 ET neurons for PV+ inputs in the mouse somatosensory cortex. Among other cell-surface molecules, *Cdh12* and *Cdh13* are differentially enriched between these two populations of pyramidal cells. Both cadherins are capable of homophilic interactions ^43,44^ and show specific expression patterns within the interneuron subtype that match the preferred inhibitory profile of each pyramidal cell. Previous studies suggested that the chemokine Cxcl12 might direct PV+ axons to target L5 ET neurons ^45^. While *Cxcl12* was not identified in our RNAseq screening, its low expression in these cells might hinder its detectability.

Even though *Cdh12* and *Cdh1*3 are expressed in both populations of L5 pyramidal cells, they exhibit different functions depending on the cellular context. Indeed, our genetic manipulations demonstrate that *Cdh13* exclusively drives PV+ interneuron wiring onto L5 ET cells, but it does not play a similar role in L5 IT neurons. Conversely, *Cdh12* only drives CCK+ basket connectivity in L5 IT cells despite also being expressed in L5 ET neurons. Why is the role of *Cdh12* and *Cdh13* in instructing CCK+ and PV+ inputs not conserved across different types of L5 pyramidal cells? Cdh13 is an atypical cadherin superfamily member as it lacks a transmembrane domain and is inserted into the plasma membrane via a glycosyl-phosphatidylinositol anchor ^46^. As such, Cdh13 requires other proteins for downstream signaling ^47^ and may interact with different synaptic partners depending on the pyramidal cell type. It is conceivable that L5 IT and L5 ET neurons express different interactomes and may not contain the relevant postsynaptic machinery to allow *Cdh13* and *Cdh12* to instruct PV+ and CCK+ basket cell wiring, respectively. Identifying the co-receptors and downstream signaling mechanisms that regulate cadherin function in this cell type-specific context would shed light on this process.

Trans-synaptic rabies tracing using glycoprotein (G)-deleted rabies virus is a powerful and versatile tool to explore circuit connectivity and uncover the distribution of inputs onto specific neuronal populations ^48^. We engineered a multiplex monosynaptic tracing approach to visualize inhibitory inputs onto L5 IT and L5 ET cell types in the same brain, based on previous studies ^36,49^. Simultaneous mapping of the presynaptic networks of L5 IT and L5 ET neurons allowed us to examine whether L5 pyramidal cells received inhibition from the same or segregated subtypes of PV+ interneurons. Although our experiments revealed an unbalanced number of starter cells between the two populations of L5 pyramidal cells, we labeled a similar number of inputs for both L5 pyramidal cell types. We found that L5 ET neurons receive a larger fraction of inputs per starter cell than L5 IT cells, reinforcing that L5 ET neurons receive more inhibitory inputs than L5 IT cells and may require more inhibitory control to maintain their excitatory-inhibitory balance.

Our experiments also unveiled that L5 IT and L5 ET neurons received inhibitory inputs from largely segregated presynaptic networks. To our knowledge, such exclusive connectivity patterns have not been described before in the cerebral cortex. Previous studies instead indicate that neighboring cells share inputs ^4–6,42^. Our results suggest that such connections are rare and that L5 IT and L5 ET neurons participate in separate networks. It should be noted, however, that the incomplete mapping of all inhibitory inputs to all L5 pyramidal cells undoubtedly lowered the probability of identifying PV+ interneurons connected to both L5 IT and L5 ET neurons. Nevertheless, the different spatial distribution of L5 IT and L5 ET PV+ inputs and the diverse expression patterns of *Cdh13* among different PV+ types strongly suggest that segregated PV-to-pyramidal cell type connections are a rule of inhibitory connectivity in the cerebral cortex, at least for L5. Another possible caveat of these experiments is the minimal fraction of co-labeled inputs compared to single-labeled inputs, which may arise from the competition of distinct rabies types and jeopardize our capacity to visualize shared inputs. Although we may certainly underestimate the contribution of shared inputs, the fact that we could identify excitatory and inhibitory neurons infected with GFP and tdTomato RV confirms that co-infections are biologically possible in our system. Finally, we cannot exclude the possibility that L5 IT and L5 ET neurons share other types of inputs, although recent work demonstrates that at least some subtypes of somatostatin interneurons also target specific populations of pyramidal cells (Wu et al., 2023).

Previous studies have highlighted the existence of translaminar PV+ inhibition in L5 pyramidal cells ^50–53^. Our results revealed that L5 ET neurons receive inhibition primarly from local L5 PV+ interneurons, while L5 IT neurons receive inputs from PV+ cells spread across different layers, with similar local and translaminar inhibitory contributions. At least ten PV+ interneuron subtypes with unique transcriptomic profiles and laminar positions have been identified ^34^. We found that the expression profile of *Cdh13* in the different subtypes of PV+ interneurons mirrors the spatial distribution of PV+ inputs preferentially targeting L5 ET cells. Hence, the laminar distribution of PV+ inputs targeting L5 IT and L5 ET cells seem to reflect the existence of different inhibitory circuits organized by the expression of matching cell-surface molecules.

Extensive work has demonstrated the dichotomy of PV+ and CCK+ perisomatic inhibition. PV+ basket cells are often described as clockworks for cortical network oscillations, whereas CCK+ basket cells are seen as modulators of such activity ^22,54^. Both basket cells exhibit different intrinsic properties and fire in different phases of behavioral activity ^55^, but some studies suggest that the function of these interneurons is interlinked. Experience-dependent or experimentally-induced changes in the activity of pyramidal cells cause bidirectional changes in persisomatic inhibition in the hippocampus ^56,57^. Specifically, increased activity in pyramidal cells enhances PV-mediated inhibition and reduces CCK-mediated inhibition ^56,57^. Here, we found no interdependency between both types of perisomatic inhibition, at least at the input level. Modifying the density of one input – by interfering with *Cdh12* or *Cdh13* levels - did not impact the other, supporting the idea that PV+ and CCK+ basket cells have non-overlapping functions. Our results also suggest that different regulatory mechanisms control the development of the inhibitory wiring and the plasticity of pyramidal neurons.

Why would some neurons need both types of inhibition while others receive barely any CCK+ inhibition? The two populations of L5 pyramidal cells consistently exhibit distinct functions across the different cortical areas. For example, L5 ET neurons exhibit higher contrast sensitivity and broader tuning response than L5 IT neurons ^58^. During the early phases of a task, L5 IT cells are actively involved, whereas L5 ET neurons become increasingly recruited in the sensory, motor, and prefrontal cortices as the task progresses ^1,59,60^. Evidence also supports the notion that L5 IT neurons uphold and enhance the computational efficiency of all cortical neurons by continuously updating them. In contrast, L5 ET neurons serve as information broadcasters, relaying the updated information to subcortical structures ^61^. Our work shows that each population of L5 pyramidal cells receives a different combination of perisomatic inputs. L5 ET cells receive dense and almost exclusively PV+ innervation, whereas both PV+ and CCK+ cells contribute to the perisomatic inhibition of L5 IT cells. Since L5 ET neurons send their outputs outside the cortex, these cells might need tighter regulation. Indeed, mice demonstrate a more adept ability to regulate the activity of ET neurons compared to IT neurons ^62^. In contrast, L5 IT cells might compute more diverse information and require increased plasticity ^62^ CCK+ inputs may confer more precision in the modulation of L5 IT neurons, a phenomenon further reinforced by modulatory pathways ^63,64^.

Considering the unique function of L5 pyramidal cells, it is foreseeable that any reduction or mistargeting of their inhibitory inputs could significantly disrupt the circuitry, and lead to various pathological conditions ^65^. For instance, L5 pyramidal cells seem particularly vulnerable in autism ^66,67^. Future investigations to understand the relationship between synaptic excitatory/inhibitory ratio within cortical circuits, input specificity and risk gene expression might shed some light on the etiology of some neurodevelopmental disorders.

## Author contributions

J.J. and B.R. conceived and designed the study. J.J. performed most of the experiments and analyzed all data. G.C. designed and produced the molecular tools, with the help of T.G. and P.M. G.C. and T.G. conducted the rabies production. T.G. and P.M. helped with the *in vitro* experiments and the AAV virus production. S.S., T.G., P.M. helped with the tissue preparation and immunohistochemistry. T.K. also contributed to collecting and analyzing data on the synaptic developmental timeline. S.S. performed the Cdh12/13 KD *in situ* validation and analyzed the data. F.H. performed the RNA-Seq data analysis and visualization. T.K. and M.B. performed the *ex-vivo* recordings and analyzed the data. J.J. and B.R. wrote the manuscript with inputs from all authors.

## Acknowledgements

We are thankful to L. Doglio, F. Sanchez-Roman, S. Sanalidou and T. Garces for technical assistance and lab support, and to Ian Andrew for mouse management. The pAAV *hSyn-fDIO-MCS* construct was kindly shared by M. Selten and O. Marín (King’s College London) The *pSAD ΔG F3* construct was kindly provided by M. Tripodi (University of Cambridge). B7GG and HEK293-TVA cells were generously shared by T. Karayannis (University of Zurich). The *pSAD ΔG ChR2c::T2A::tdTomato* construct and the *RVdG-EnvA-tdTomato-ChR2* viral preparation were generously shared by K. Conzelmann (Ludwig-Maximilians-University Munich) and A. Delogu (KCL). We thank the IoPPN Genomics & Biomarker Core Facility, the Advanced Cytometry Platform of the Research and Development Department at Guy’s and St Thomas’ NHS Foundation Trust for their technical advice and assistance on the FACS experiment, and the CRG Genomics Core Facility of Barcelona for conducting RNA sequencing. Finally, we are grateful to O. Marín, J. Burrone, M. Grubb and C. Bernard for critical reading of the manuscript, and to all members of the Rico and Marín laboratories for stimulating discussions and ideas. This project was supported by grants from Wellcome Trust (202758/Z/16/Z) and European Union’s Horizon 2020 research and innovation program (AIMS- 2- TRIALS, 777394) to B.R, and the EMBO Long-Term Fellowship to J.J. and G.C.

---

**Extended Data Fig. 1.**
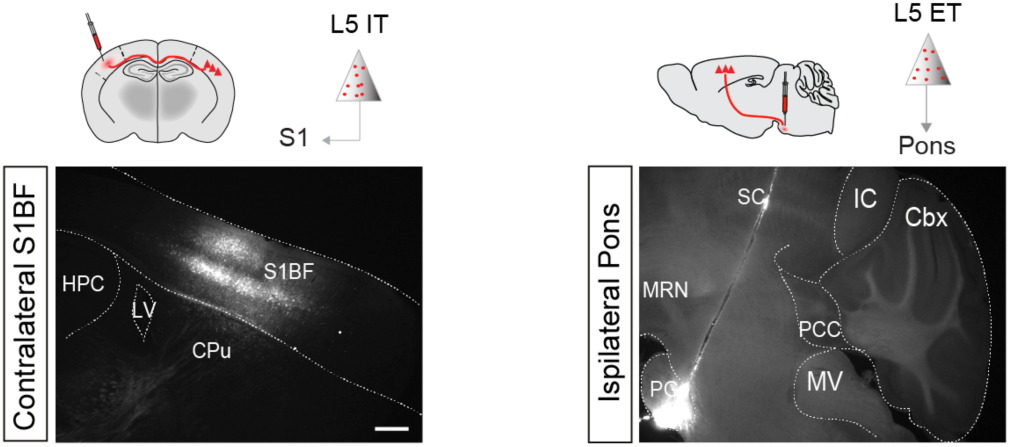
Retrobeads injection site. Confocal images of retrobeads injection site in the barrel field area of the somatosensory cortex and in the pons to target L5 IT and L5 ET neurons, respectively. Scale bar, 500μm

**Extended Data Fig. 2.**
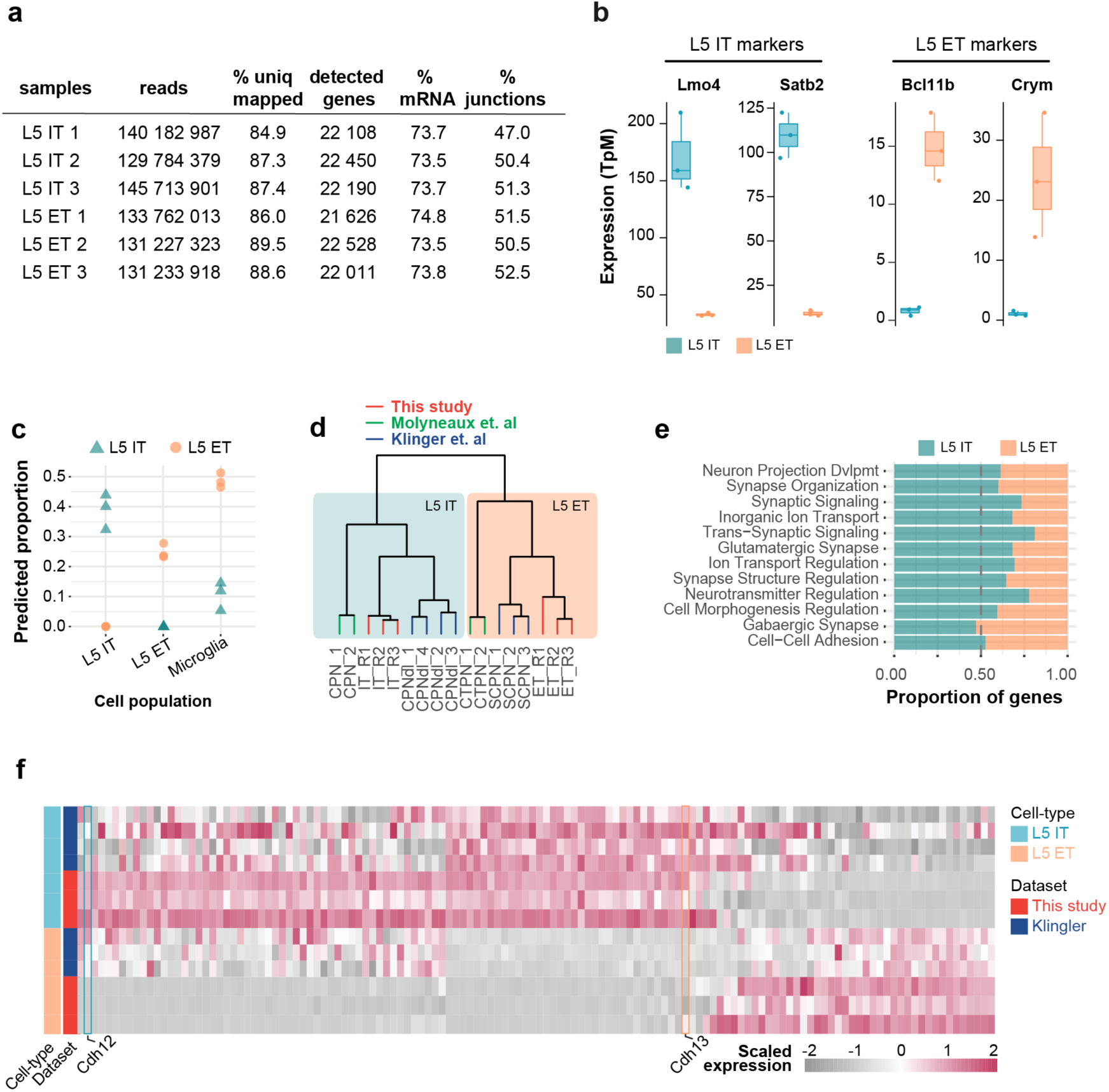
RNA-sequencing data. **a,** FastQC analysis of RNA sequencing data for each experimental replicate showing the total number of reads and the percentage of reads uniquely mapped to the reference genome, number of detected genes, mRNA representation and reads mapped to exon-exon junctions. **b,** Expression levels (TpM) of key markers genes in L5 IT and L5 ET pyramidal cell types. **c,** Estimated cell type proportions (L5 IT, L5 ET and microglia) after deconvolution of our RNAseq dataset. L5 IT replicates are denoted as triangles, while L5 ET replicates are denoted as dots. **d,** Dendrogram of this study, Molyneaux et al., and Klinger et al., with branches annotated by experimental replicate and L5 pyramidal cell types. Note the low distance among replicates and how samples cluster by L5 pyramidal cell type. **e,** Proportion of gene ontology terms enriched in L5 IT or L5 ET cell types. **f,** Heatmap showing DEG between L5 IT and L5 ET neurons in this study and Klingler et al.

**Extended Data Fig. 3.**
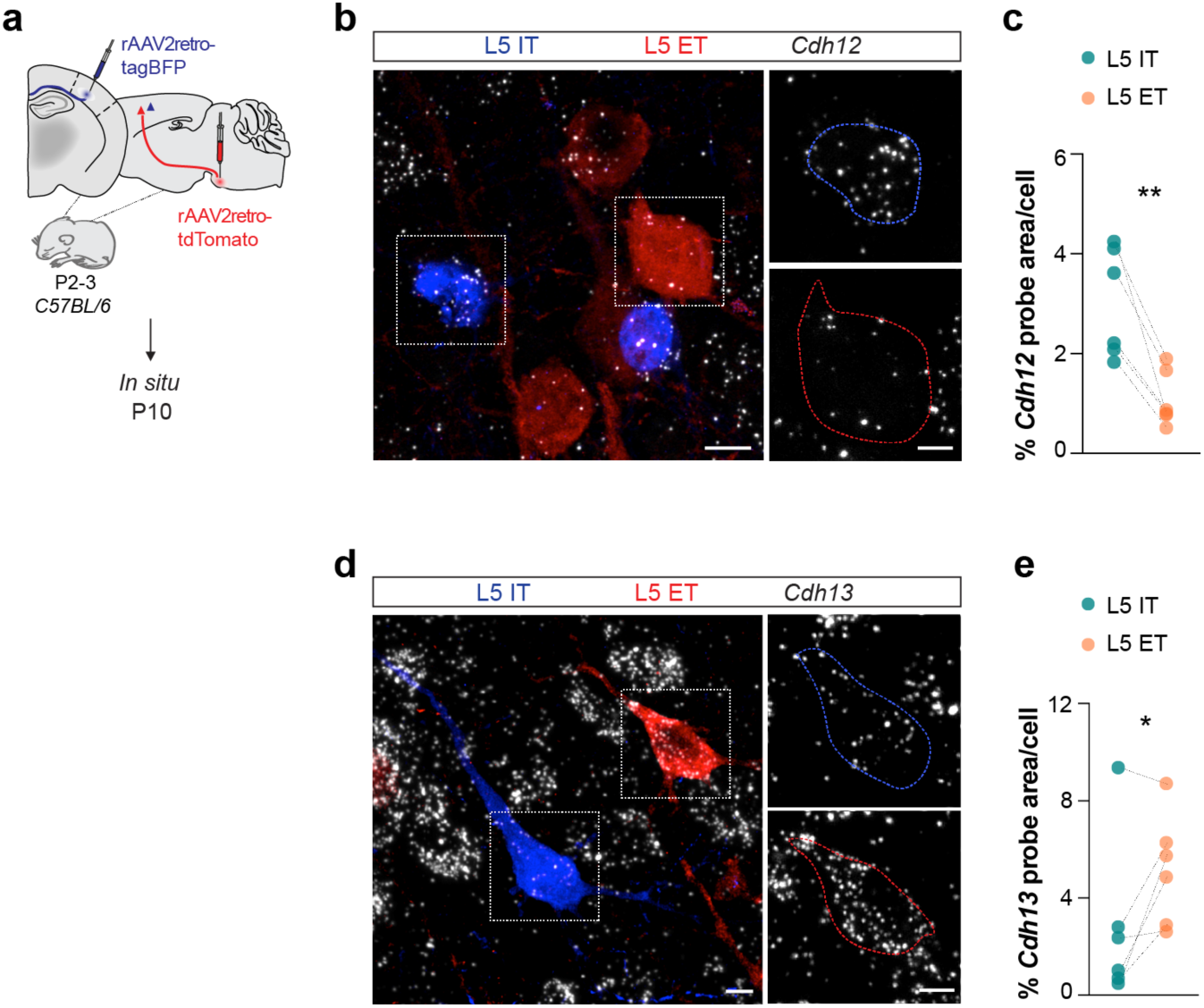
*In situ* validation of candidate genes enrichment in L5 IT and L5 ET neurons. **a,** Candidate genes mRNA expression levels in L5 IT and L5 ET neurons assessed by single-molecule fluorescent i*n situ* hybridization at P10. **b,** Confocal images illustrating *Cdh12* RNA expression in retrogradely-labeled L5 IT (blue) and L5 ET (red) neurons. **c,** Fraction of *Cdh12* probe area normalized per cell area (*L5 IT* and *L5 ET* n = 6 mice; paired t-test **P < 0.01). **d,** Confocal images illustrating *Cdh13* RNA expression in retrogradely-labeled L5 IT (blue) and L5 ET (red) neurons. **e,** Fraction of *Cdh13* probe area normalized per cell area (*L5 IT* and *L5 ET* n = 6 mice; paired t-test \**P* < 0.05). Data are represented as mean ± s.e.m. Scale bars, 5 μm, 10 μm (b, d).

**Extended data Fig. 4.**
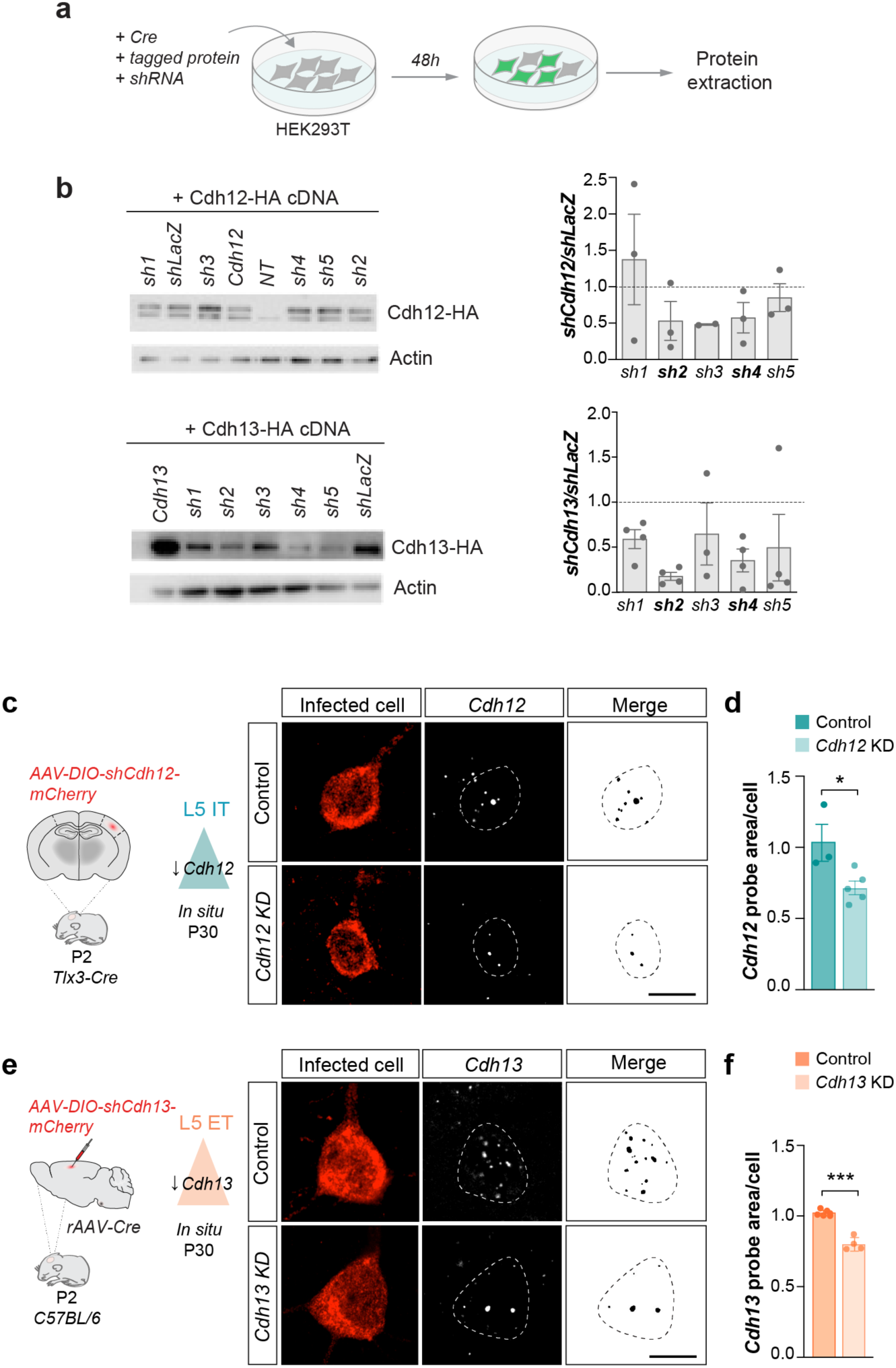
*In vitro* selection of shRNA and *in vivo* validation of *Cdh12* and *Cdh13* down-regulation. **a,** Experimental design. **b,** Left: *in vitro* protein expression assessed by Western blot of *Cdh12* and *Cdh13* HA-tagged constructs from transfected HEK923T cells. Right: Quantification of protein signal normalized to actin for each *shRNA*, relative to control transfections with a *shLacZ*-expressing plasmid (n = 3 independent cultures). **c,** Representative confocal images of *Cdh12* RNA expression (white) at P30 in L5 IT cells infected with control *shLacZ-* or *shCdh12*-mCherry (red). **d,** Normalized *Cdh12* RNA expression in L5 IT (*Control* n = 3 mice; *Cdh12 KD* n = 5 mice; one-tailed t-test \**P* < 0.05). (E) Representative confocal images of *Cdh13* RNA expression (white) at P30 in L5 ET cells infected with control *shLacZ*- or *shCdh13*-mCherry (red). **f,** Normalized *Cdh13 RNA* expression in L5 ET (*Control* n = 5 mice, *Cdh13 KD* n = 4 mice, one-tailed t-test \*\*\**P* < 0.001). Data are represented as mean ± s.e.m. \**P* < 0.05; \*\**P* < 0.01; \*\*\**P* < 0.001; ns, not significant. Each dot represents an individual mouse. Data are mean ± s.e.m. Scale bar, 10μm (c, d).

**Extended data Fig. 5.**
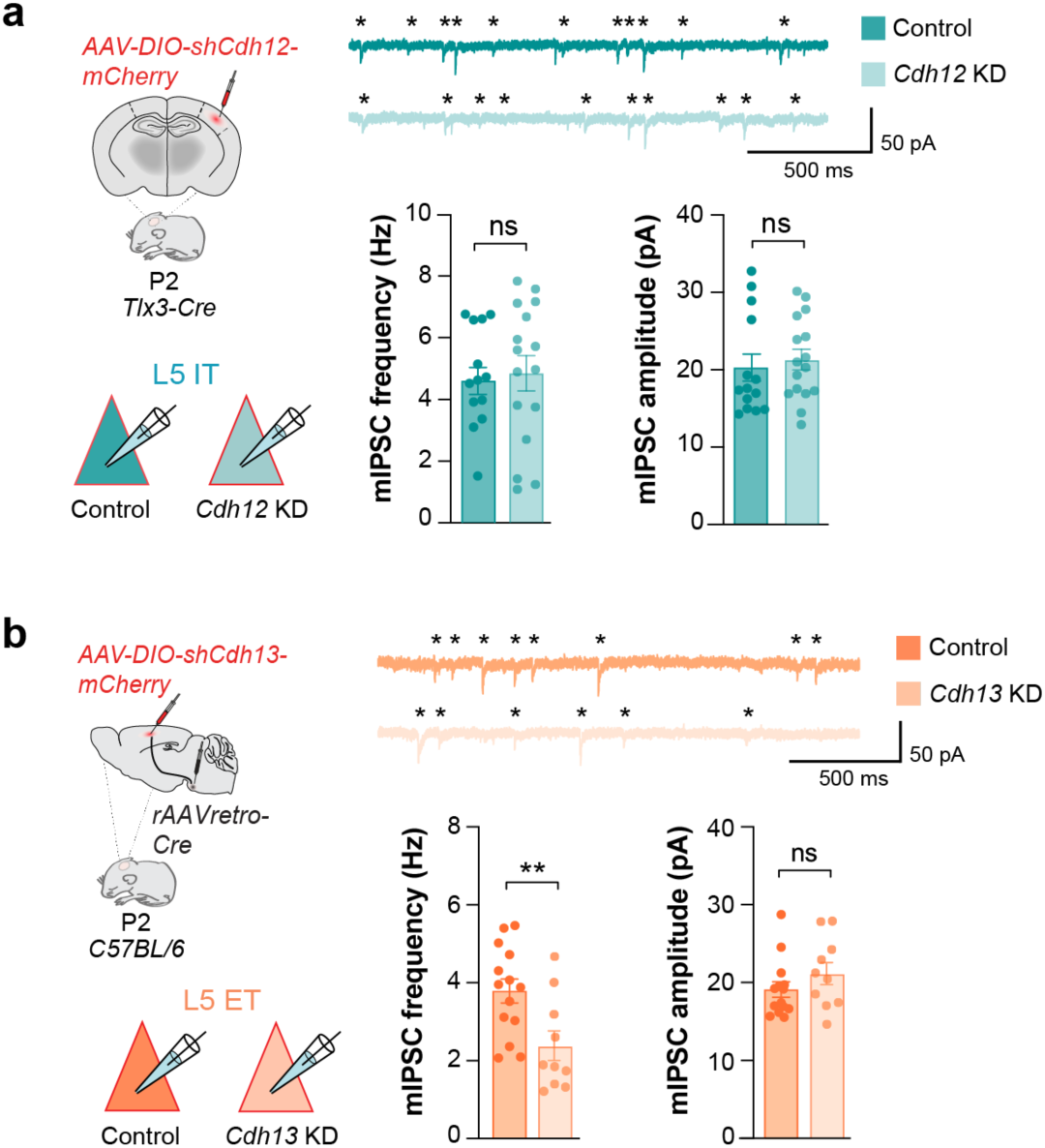
Functional synaptic outcome of *Cdh12* KD in L5 IT neurons and *Cdh13* KD in L5 ET neurons. **a,** Left: Experimental design to selectively down-regulate *Cdh12* in L5 IT and record from control or *Cdh12* KD infected cells. Top: Example traces of mIPSCs recorded from L5 IT cells infected with a control *shLacZ* or *shCdh12*. Bottom: Quantification of the frequency (left) and amplitude (right) of mIPSCs from L5 IT cells in both conditions (*Control* n = 14 cells, 5 mice, *Cdh12 KD* n = 16 cells, 5 mice; t-test frequency and amplitude ns). **b,** Left: Experimental design to selectively down-regulate *Cdh13* in L5 ET and record from control or *Cdh13* KD infected cells. Top: Example traces of mIPSCs recorded from L5 ET cells infected with a control *shLacZ* or *shCdh13*. Bottom: Quantification of the frequency (left) and amplitude (right) of mIPSCs from L5 ET cells in both conditions (*Control* n = 14 cells, 4 mice, *Cdh13 KD* n = 10 cells, 4 mice, t-test frequency \*\**P* < 0.01, amplitude Mann-Whitney test ns). Data are represented as mean ± s.e.m.

**Extended data Fig. 6.**
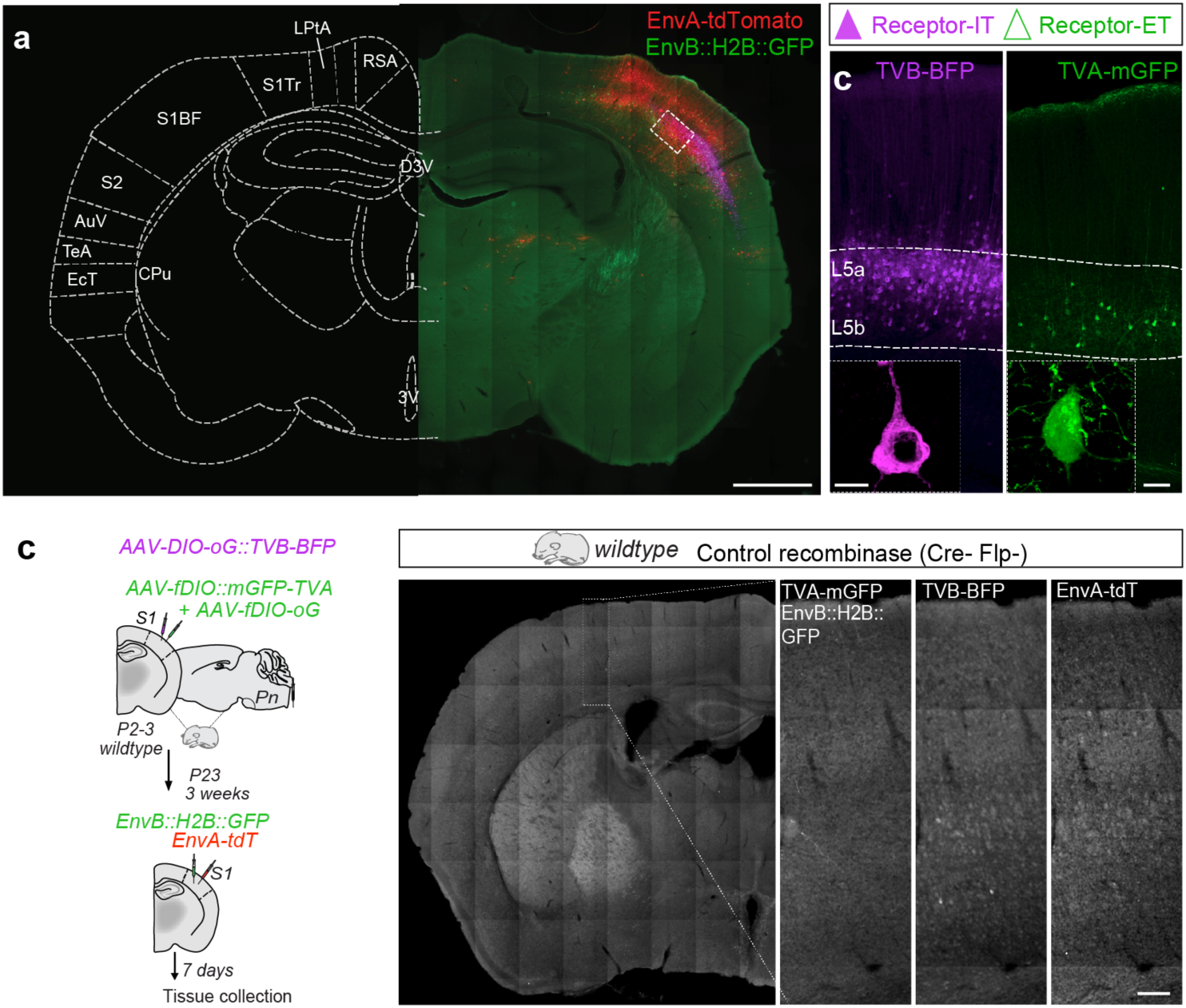
Cell type-specific expression of the multiplex monosynaptic tracing strategy. **a,** Coronal section depicting TVA/TVB-expressing AAVs and EnvA/EnvB-expressing RV injection sites. **b,** Confocal images illustrating the distribution of L5 IT and L5 ET receptor-expressing cells. L5 IT neurons expressing TVB-BFP (magenta) were mainly present in L5a, while L5 ET expressing TVA-mGFP (green) were exclusively found in L5b. **c,** Confocal images illustrating the absence of receptor- or RV-infected cells in a Cre- and Flp-negative brain. Scale bars, 500μm (a) 50 μm (b, c), 10 μm (b).

## Methods

### Animals

*Tlx3^Cre^* ^33^ and *Nex^Cre^* ^68^ were maintained in a C57BL/6 background (Charles River Laboratories). Animals were housed in groups of up to five littermates and maintained under standard, temperature controlled, laboratory conditions. Mice were kept on a 12:12 light/dark cycle and received water and food *ad libitum*. All animal procedures were approved by the ethical committee (King’s College London) and conducted in accordance with European regulations, and Home Office personal and project licenses under the UK Animals (Scientific Procedures) 1986 Act.

### Plasmids design

#### shRNA

*Cdh12* and *Cdh13 shRNA* were cloned into a *pAAV-EF1a-DIO-mCherry* vector as previously described ^32^. The ssDNA primers to generate the *shRNAs* were obtained using the Block-it RNAi web tool (Thermo Scientific) and were as follows: *shCdh12#1* (Fwd: CTA GGC ATT CGG ACT TGG ATA AAG GCC TGA CCC ACC TTT ATC CAA GTC CGA ATG CTT TTTG and Rev: AAT TCA AAA AGC ATT CGG ACT TGG ATA AAG GTG GGT CAG GCC TTT ATC CAA GTC CGA ATGC); *shCdh12#2* (Fwd: CTA GGC AGT ACC AGG TCC TCA TTC ACC TGA CCC ATG AAT GAG GAC CTG GTA CTG CTT TTTG and Rev: AAT TCA AAA AGC AGT ACC AGG TCC TCA TTC ATG GGT CAG GTG AAT GAG GAC CTG GTA CTGC; *shCdh12#3* (Fwd: CTA GGC TGG GCC ATT TAA GGA TAC TCC TGA CCC AAG TAT CCT TAA ATG GCC CAG CTT TTTG and Rev AAT TCA AAA AGC TGG GCC ATT TAA GGA TAC TTG GGT CAG GAG TAT CCT TAA ATG GCC CAGC); *shCdh12#4* (Fwd: CTA GGC AAT TCT CCT TTA GAT TAG CCC TGA CCC AGC TAA TCT AAA GGA GAA TTG CTT TTTG and Rev: AAT TCA AAA AGC AAT TCT CCT TTA GAT TAG CTG GGT CAG GGC TAA TCT AAA GGA GAA TTGC); *shCdh12#5 (*Fwd: CTA GGC ACG AAT ACA ATG ACTAT TCC CTG ACC CAG AAT AGT CAT TGT ATT CGT GCT TTT TG and Rev: AAT TCA AAA AGC ACG AAT ACA ATG ACT ATT CTG GGT CAG GGA ATA GTC ATT GTA TTC GTGC); *shCdh13#*1 (Fwd: CTA GGC TCC TTG CAG GAT ATC TTT ACC TGA CCC ATA AAG ATA TCC TGC AAG GAG CTT TTTG and AAT TCA AAA AGC TCC TTG CAG GAT ATC TTT ATG GGT CAG GTA AAG ATA TCC TGC AAG GAGC); *shCdh13#2* (Fwd: CTA GGG GCT GCA TAC ACC ATC ATC ACC TGA CCC ATG ATG ATG GTG TAT GCA GCC CTT TTTG and Rev: AAT TCA AAA AGG GCT GCA TAC ACC ATC ATC ATG GGT CAG GTG ATG ATG GTG TAT GCA GCCC); *shCdh13#3* (Fwd: CTA GGC TGA TCA AAG TGG AGA ATG ACC TGA CCC ATC ATT CTC CAC TTT GAT CAG CTT TTTG and Rev: AAT TCA AAA AGC TGA TCA AAG TGG AGA ATG ATG GGT CAG GTC ATT CTC CAC TTT GAT CAGC); *shCdh13#4* (Fwd: CTA GGC CTT CTT CAG AAT CTG AAC ACC TGA CCC ATG TTC AGA TTC TGA AGA AGG CTT TTTG and Rev: AAT TCA AAA AGC CTT CTT CAG AAT CTG AAC ATG GGT CAG GTG TTC AGA TTC TGA AGA AGGC); *shCdh13#5* (Fwd: CTA GGC CCA TCA TGG TGA CAG ATT CCC TGA CCC AGA ATC TGT CAC CAT GAT GGG CTT TTTG and Rev: AAT TCA AAA AGC CCA TCA TGG TGA CAG ATT CTG GGT CAG GGA ATC TGT CAC CAT GAT GGGC). The same *shLacZ* as described in ^32^ was used as a control condition.

#### Rabies

Constructs used for the multiplex G-deleted rabies virus-mediated circuit mapping approach were generated by standard molecular biology cloning procedures. The *pSAD 11.G H2B:GFP* was obtained by taking advantage of the *pSAD 11.G F3* expression vector (kindly provided by M. Tripodi, Cambridge University, AddGene #32634). The nucleotide sequence encoding the histone H2B-tagged GFP was designed and ordered via GeneArts (ThermoFisher Scientific). Subsequently, it was subcloned within the *pSAD 11.G F3* expression vector by a restriction enzyme-based method, using the unique restriction enzymes AscII and BsiWI (New England Bioscience). The membrane-bound *EGFP::T2A::TVA* cassette was obtained by gene synthesis. A DNA fragment containing the nucleotide sequence of the membrane-bound EGFP link via a T2A cleaving site to the TVATC66 receptor Field 69 coding sequence was designed in an inverted orientation and ordered via GeneArts (ThermoFisher Scientific). This DNA sequence was subsequently amplified by Q5 Hot Start High Fidelity DNA polymerase-based PCR (New England Bioscience) using primers containing the unique restriction sites AccI and NheI (Fwd: TAA GCA <u>GTC GAC</u> TTA CTT GGA TGC GCT TTC AAG and Rev: TGC TTA <u>GCT AGC</u> GCC ACC ATG CTG TGC TGT ATG). The amplified sequence was then inserted within the expression vector *pAAV-hSyn-fDIO-MCS* (kindly provided by Dr. M. Selten and Dr. O. Marín) using the AccI and NheI sites.

The same experimental approach has been used to generate the *pAAV-hSyn fDIO oG* expression vector. The oG nucleotide sequence was PCR amplified using specific primers containing the restriction sites AccI and NheI (Fwd: TAA GCA <u>GTC GAC</u> TTA GAG CCG TGT CTC GCC and Rev: TGC TTA <u>GCT AGC</u> GCC ACC ATG GTC CCA CAG GCT CTC CTC) and then cloned into *pAAV-hSyn-fDIO-MCS.* The *pAAV-TVB-tagBFP::T2A::oG* was designed as previously described ^36^ and ordered via GeneArts (ThermoFisher Scientific). The cassette was then sub-cloned into the expression vector *pAAV-hSyn Flex tdTomato::T2A::SypEGFP* (AddGene #51509) using the AscI and FseI restriction sites. All constructs were verified by sequencing.

### Cell culture and transfection

HEK293T cells were cultured in Dulbecco’s Modified Eagle’s medium supplemented with 10% fetal bovine serum, 2 mM glutamine, penicillin (50 units/ml) and streptomycin (50 g/ml). The cultures were incubated at 37°C in a humidified atmosphere containing 5% CO_2_. HEK293T cells were transfected using polyethylenimine (PEI, Sigma) at a 1:4 DNA:PEI ratio or Lipofectamine 2000 (Thermo Fisher Scientific).

### AAV and rAAVretro production

HEK293FT cells (Thermo Fisher Scientific R70007) were seeded on 15-cm plates and co-transfected with packaging plasmids AAV-ITR-2 genomic vectors (7.5 µg), AAV-Cap8 vector pDP8 (30 µg; PlasmidFactory GmbH, Germany, #pF478) or AAV-Cap DJ Rep-Cap vector (30 µg; Cell Biolabs, VPK-420-DJ) using PEI (Sigma) at a ratio 1:4 (DNA:PEI). 72 hours post-transfection, supernatants were incubated with Ammonium sulfate (65g/200ml supernatant) for 30 minutes on ice and centrifuged for 45 minutes at 4000 RPM at 4°C. Transfected cells were harvested and lysed (150mM NaCl, 50mM Tris pH8.5), followed by three freeze-thaw cycles and Benzonase treatment (50U/ml; Sigma E1014-25KU) for 1 hour at 37°C. Filtered AAVs (0.8 µm and 0.45 µm MCE filters) from supernatants and lysates were run on an Iodixanol gradient by ultracentrifugation (Vti50 rotor, Beckmann Coultier) at 50,000 RPM for 1 hour at 12°C. The 40% iodixanol fraction containing the AAVs was collected, concentrated using 100 kDa-MWCO Centricon plus-20 and Centricon plus-2 (Merck-Millipore), aliquoted and stored at -80°C. The number of genomic copies was determined by qPCR using the following primers against the WPRE (Fwd: GGC ACT GAC AAT TCC GTG GT and Rev: CGC TGG ATT GAG GGC CGAA). AAVs with a titer equal or higher to 10^12^ genome copy/ml were used for *in vivo* injections. For down-regulation experiments, the two shRNAs with the most efficient down-regulation *in vitro* were used for each target gene. The two shRNA plasmids were co-transfected for AAV production in order to increase the down-regulation efficiency.

### Rabies production

#### Cell lines

HEK-TVA (The Salk Institute of Biological Sciences), HEK-TVB (The Salk Institute of Biological Sciences), BHK-EnvA (kindly gift from T. Karayannis, University of Zurich), BHK-EnvB (The Salk Institute of Biological Sciences), and B7GG (kind gift from T. Karayannis) cells were maintained in DMEM (Gibco), supplemented with 10% fetal bovine serum (FBS) in a humified atmosphere of 3% CO_2_ and 35°C.

#### G-deleted rabies virus production

G-deleted rabies viruses (RV11.G) were produced as previously described ^49^. Briefly, *RV11.G-H2B:GFP* and *RV11.G-tdTomato* were recovered in B7GG cells by Lipofectamine 2000 (Thermo Fisher Scientific) transfection with *pcDNA-SADB19N*, *pcDNA-SADB19P*, *pcDNA-SADB19L* and *pSAD11.G H2B:GF*P or *pSAD11.G:tdTomato* (kind gift from K. Conzelmann). During the virus production, the transfected cells were maintained using DMEM medium (Gibco) supplemented with 10% FBS in a humified atmosphere of 3% CO_2_ and 35°C.

For pseudotyping with EnvA and EnvB, BHK-EnvA and BHK-EnvB cell lines were infected with unpseudotyped *SAD 11.GtdTomato* and *SAD 11.GH2B:GFP* rabies viruses, respectively. Subsequently, infected cells were washed with PBS, collected by 0.25% trypsin-EDTA and replated in new dishes. The virus-containing medium was then filtrated via 0.45μm filters (Cornings) and concentrated through two rounds of ultra-centrifugation. The infectious titers of the purified viruses were determined using the HEK-TVA and HEK-TVB cell lines. In addition, the presence of any contamination of unpseudotyped rabies virus was detected by using HEK293T cells. Aliquots containing pseudotyped rabies viruses were stored at -80C.

### Stereotaxic Injections

#### Retrobeads

L5 pyramidal cell types were retrogradely labelled using green (488nm) or red (555nm) fluorescent retrobeads IX (Lumafluor Corp., FL). P2-3 pups were anesthetized with isoflurane and mounted on a stereotactic frame using a 3D printed isoflurane mask. Unilateral injections of 75nl retrobeads at 30nl/min were carried out as follows. L5 IT were labelled by targeting the contralateral somatosensory cortex (S1) (AP +1.6, ML -1.9 to -2.2, DV -0.8 to -0.5), L5 ET were labelled by targeting the Pons (AP -0.3, ML +0.3, DV -4.5 to -4.0). Retrobeads were undiluted and sonicated prior to each injection to avoid aggregate formation. The pipette was retracted from the brain after 2min to allow for diffusion.

#### AAV Viral injections

For i*n situ* hybridization experiments, we followed the same experimental design as previously described to target L5 IT and L5 ET cell types. Briefly, 300nl of *rAAV2retro-Ef1a-tagBFP* (1,02.10^12^ vg/ml) and 300nl of *rAAV2retro-Ef1a-NLS-tdTomato* (4,82.10^12^ vg/ml) were injected at 60nl/min in the contralateral S1 and ipsilateral Pons of P2/3 WT pups.

For *Cdh12* and *Cdh13* down-regulation experiments, we injected 300nl of *AAV8-Ef1a-DIO-shCdh12-mCherry* (8,20.1^11^ vg/ml) or *AAV8-Ef1a-DIO-shCdh13-mCherry* (9,1.10^11^ vg/ml) or *AAV8-DIO-shLacZ-mCherry* (1,20.10^12^ vg/ml) in the S1 of P2/3 pups. *Tlx3-Cre* mice were used to specifically access L5 IT, while L5 ET were specifically targeted by injecting 300nl of *rAAV2retro-Cre* (AddGene #55636-AAVrg, 2.10^13^ vg/ml) in the Pons. The stereotaxic coordinates for the different target areas were the same as for the retrobeads experiment.

#### Rabies stereotaxic injections

P2-3 *Tlx3-Cre* pups were anesthetized with isoflurane and mounted on a stereotactic frame using a 3D-printed isoflurane mask. A unilateral injection of 300nl *rAAV2retro-FlpO* (AddGene #55637-AAVrg, 1,6.10^13^ vg/ml) was made in the Pons (AP -0.3, ML +0.3, DV - 4.4 to -4.0) to target L5 ET axons. 200nl of a mix of *AAV8-DIO-oG* (1,65.10^12^ vg/ml) and *AAV8-DIO-hSyn-TVB-BFP* (4,37.10^13^ vg/ml) (2:1 mix ratio), and 200nl of a mix of AAV8-FRT-oG (9,61.10^13^ vg/ml) and *AAV8-FRT-TVA-mGFP* (9,87.10^13^ vg/ml, 2:1 mix ratio) were co-injected in the ipsilateral somatosensory cortex (S1, AP +1.6, ML -1.9 to -2.2, DV -0.8 to -0.5). All viral injections were made at a 60nl/min rate using a Micro2T nanoinjector (WPI). The pipette was retracted from the brain after 5min, and the wound was closed using VetBond glue. Three weeks later, a second injection of 500nl of a mix of *RV11.G-EnvB-H2B:GFP* (1,6.10^10^ TU/ml) and *RV11.G-EnvA-tdTomato* (3,65.10^8^ TU/ml, kind gift from A. Delogu) (1:1 ratio) was made in the ipsilateral S1BF (AP-1.8, ML 3.2, DV -0.7 --> -0.4). After suturing and disinfecting with Betadine, mice received a subcutaneous injection of Buprenorphine (0.03 mg/ml) to prevent acute pain. Note that the stereotaxic coordinates for the different target areas were determined from the lambda in pups (Atlas of the Developing Mouse Brain ^70^) and the bregma in young adults ^71^.

### Tissue dissociation and Fluorescence-Activated Cell Sorting (FACS)

P2-3 C57BL/6 mice were injected, as previously described, with Red Retrobeads™ IX (RB555, LumaFluor) either in S1BF to target L5 IT or in the pons to label L5 ET. To isolate individual cells, we euthanized mice at P10, extracted the brain and microdissected lower layers of S1 in cold pH 7.3 dissociation media containing 14mM MgCl_2_, 2mM HEPES (Invitrogen 15630-106), 0.2mM NaOH (Sigma S0899), 90mM Na_2_SO_4_ (Sigma S6547), 30mM K_2_SO_4_ (Sigma P9458), 3.6mg/mL D-(+)-Glucose (Sigma G6152), 0.8mM kynurenic acid (Sigma K3375), 50 µM AP-V (Sigma A5282), 50U/mL penicillin/streptomycin (Thermo Fisher 15140122). To generate single-cell suspensions, 1 mm^3^ tissue pieces from 2-3 brains were pooled and enzymatically digested in a dissociation medium containing 0.16 mg/mL cysteine (Sigma C9768), 7 U/mL Papain (Sigma P3125), 0.1mg/mL DNase (Sigma 10104159001) at 37°C for 30min. Papain digestion was then blocked with a dissociation medium containing 0.1mg/mL ovomucoid (Sigma, St. Louis, MO T2011) and 0.1mg/mL bovine serum albumin (BSA, Sigma A4161) for 1 min at room temperature. Neurons were mechanically dissociated to create a single cell suspension in iced OptiMEM solution containing 3.6mg/mL D-(+)-Glucose, 4 mM MgCl_2_, 0.4mM kynurenic acid, 25 µM AP-V, 0.04 mg/mL DNase, diluted in OptiMEM medium (Thermo Fisher 31985). Actinomycin D (Sigma A1410) was added during the dissociation process to protect the tissue and prevent activation of immediate early genes ^72^. Cells were centrifugated at 120g for 5 min at 4°C, resuspended in 150-300 µL of fresh complemented OptiMEM and passed through a 40 µm cell strainer. Retrobeads-labelled individual cells were then purified from the cell suspension using a BD FACS Aria III cytometer. Cells were collected in 350μl of RLT buffer (RNeasy Lysis buffer, QIAGEN) containing 1% 2-Mercaptoethanol and stored at -80°C for RNA extraction.

### RNA-sequencing

RNA was extracted using the QIAGEN RNeasy Micro Kit according to the manufacturer’s instructions. Library preparation and RNA-seq experiments were performed by the Genomic Unit of the Centre for Genomic Regulation (CRG, Barcelona, Spain). Approximately 10,000-20,000 cells were required to obtain 5-10 ng of total RNA, which served as input for the library preparation using the SMARTer v4 Ultra Low RNA Kit. The Illumina HiSeq 2500 platform was used to sequence libraries to a mean of approximately 100 million mapped 125 base pair paired-end reads per sample. In the RNA-seq experiments, three biological replicates were ascertained for each dataset.

### RNA-sequencing data processing and differential expression analysis

Our dataset (this study) was enriched with L5-specific genes using two other complementary RNA-seq datasets: L5-specific genes at P10 ^26^ and L5-specific genes at P0 ^27^. We used the first dataset as a positive control for L5 cell type-specific gene expression at P10 ^26^. The second dataset ^27^ provides an early gene expression and allowed us to ascertain the “L5 neuronal signature” further. Sequencing files from this study and ^27^ were QC’ed, genome aligned, and gene expression quantified using Nextflow’s RNA-seq pipeline (v3.2). Ensembl’s GRCm38 annotation and genomic sequences were used as reference. Gene expression counts data were kindly provided by D. Jabaudon (Geneva University) from ^26^, and an unpublished dataset of ET cortical spinal cord pyramidal cells. Genes with at least 10 read counts in each sample replicated across all studies were kept. Differential gene expression analysis was performed to identify differentially expressed genes (DEGs) between L5 IT and L5 ET neurons in each study using the R/Bioconductor package DESeq2. Genes that showed a 1.5-fold change in expression and passed the 0.05 Bonferroni-adjusted p-values were labelled as differentially expressed. We selected DEGs from this study that are concordantly regulated in ^27^. A similar selection process was performed on DEGs identified from ^26^, and the union of genes from both datasets were used for downstream analyses.

### Gene Ontology analysis

Functional enrichment in gene functions was determined using the R/Bioconductor package ClusterProfiler. The org.Mm.eg.db package containing Gene Ontology terms was used as the database, and a list of stably expressed genes was used as background reference. Terms that passed the 0.05 FDR threshold were annotated as significantly enriched.

### Scoring of cell-adhesion molecules

A list of ligand-receptor pairs of cell-surface molecules was manually compiled from previous studies ^28–30,73^ and from the STRING database. Fold-change score of the expression of L5 IT- and L5 ET-specific adhesion molecules were calculated from the expression values from this and ^26^ studies. Essentially, the fold-difference in mean normalized gene expression between L5 IT and L5 ET cell types was calculated and scaled to produce values between 0 and 1. The specificity of the interaction between L5 IT with CCK+ interneurons, and between L5 ET with PV+ interneurons was determined by calculating the enrichment in the gene expression of partner adhesion molecules amongst interneuron subpopulations. Gene expression profiles of single cells from ^74^ were used for this part of the analysis. CCK+ interneurons were identified in the Mouse Atlas by a high expression of the *Cck* gene and moderate/low expression of the *Pvalb* gene. Essentially, log2 fold-difference in the expression of each receptor in CCK+ or PV+ interneurons was calculated by comparing its expression to other interneuron subpopulations. Receptor genes that are positively enriched in CCK+ or PV+ cells were retained and log2 fold-difference values from each ligand-receptor pair were averaged to give a representative specificity score.

### Immunohistochemistry

Animals were deeply anesthetized with sodium pentobarbital by intraperitoneal injection and then transcardially perfused with PBS followed by 4% paraformaldehyde (PFA) in PBS. Dissected brains were post-fixed for 2 h at 4°C, cryoprotected successively in 15% and 30% sucrose (Sigma S0389) in PBS, and finally cut frozen on a sliding microtome (Leica SM2010 R) at 40 µm. Free-floating brain slices were permeabilized with 0.25% Triton X-100 (Sigma T8787) in PBS for 1 h at room temperature (RT) and blocked for 2 h in a solution containing 0.3% Triton X-100, 1% serum, and 5% bovine serum albumin (BSA) (Sigma A8806) at RT. Brain slices were then incubated overnight at 4°C with primary antibodies. The next day, the tissue was repeatedly rinsed in PBS and incubated with secondary antibodies for 2 h at RT. All primary and secondary antibodies were diluted in 0.3% Triton X-100, 1% serum and 2% BSA.

For perisomatic inputs quantification, the following primary and secondary antibodies were used: mouse anti-NeuN (1:500, Sigma MAB377), rabbit anti-NeuN (1:500, Millipore ABN78), goat anti-CB1R (1:400, Frontier Institute CB1-Go-Af450-1), mouse anti-CB1R (1:500, SySy 258011), rabbit anti-DsRed (1:500, Clontech 632496), mouse anti-Syt2 (1:125, ZFIN ZDB-ATB-081002-25), chicken anti-GFP (1:1000, Aves Lab GFP-1020), rabbit anti-GFP (1:500, Molecular Probes A11122), donkey anti-rabbit 405 (1:250, Abcam Ab175652), donkey anti-goat 488 (1:500, Molecular Probes A11055), goat anti-mouse IgG2b 555 (1:500, Molecular Probes A21147), donkey anti-mouse 647 (1:500, Molecular Probes A31571) and goat anti-mouse IgG2a 647 (1:500, Molecular Probes A21241).

For mapping of L5 IT and L5 ET monosynaptic inputs, every 3 sections containing somatosensory cortex were used (8 sections per mouse) and incubated with the following primary and secondary antibodies: guinea pig anti-RFP (1:500, SySy 390004), rabbit anti-tagRFP (1:250, Evrogen AB233), chicken anti-PV (1:250, SySy 195006), donkey anti-chicken 405 (1:200, Jackson 703-475-155), goat anti-guinea pig 555 (1:500, Molecular Probes A21435) and donkey anti-rabbit 647 (1:500, Molecular Probes A31573).

### Single-molecule fluorescence in situ hybridization

P10 WT mice co-injected with *rAAV2-retro-tagBFP* and *rAAv2retro-tdTomato* were perfused as previously described. Brains were post-fixed overnight in 4% PFA in PBS, followed by cryoprotection in 30% sucrose-RNase free PBS, and finally sectioned frozen on a sliding microtome at 30 µm. For down-regulation and overexpression experiments, 40 µm slices from P30 mice were used. Fluorescent *in situ* hybridization on brain slices was performed according to manufacturer’s protocol (ACD Bio, RNAscope Multiplex Fluorescent Assay v2, #323110). *Cdh12* (ACD Bio, #842531) and *Cdh13* (ACD Bio, #443251-C3) probes were used and visualized with Opal 520 (Akoya BioScience, FP1487001KT) and Opal 650 (Akoya BioScience, FP1496001KT). Single-molecule dual-color fluorescent *in situ* hybridization was combined with immunohistochemistry, and the following primary and secondary antibodies were used: chicken anti-GFP (1:100, Aves Lab GFP-1020), rabbit anti-tagRFP (1:100, Evrogen AB233), goat anti-chicken biotin (1:200, Vector BA-9010), Streptavidin 555 (1:400, Molecular Probes S32355), goat anti-chicken 488 (1:600, Molecular Probes A11039), goat anti-chicken 568 (1:500, ThermoFisher A11041) and donkey anti-rabbit 405 (1:200, Abcam Ab175652).

### Image acquisition and analysis

For the perisomatic inputs analysis, z-stacks (100 X oil immersion objective, 1.44 NA, 2.2 digital zoom, 0.2 µm step size) of 40 µm brain slices were imaged on an inverted Leica TCS-SP8 confocal maintaining same laser power, photomultiplier gain, pinhole, and detection filter settings (1024×1024, 8 bits) and analyzed with IMARIS 7.5.2 software. The density of perisomatic PV+ and CCK+ inputs was then quantified as follows: 1) all z-stacks were submitted to a background subtraction (13.2 µm) and a Gaussian filter (0.0517 µm) step before analysis; 2) L5 IT and L5 ET NeuN+ somatas positive for retrobeads or viral infection were 3D-reconstructed using the “Create surface” tool; 3) Syt2+ and CB1R+ boutons were detected using the “Spots” tool with a spot diameter (XY filter) of 0.8 and 1.2 µm, respectively; 4) the number of Syt2+ perisomatic boutons was quantified using the “Find spots close to surface” tool (ImarisXT extension) with a 0.4 µm threshold distance applied from the 3D-reconstructed NeuN+ surface. 5) the number of CB1R+ perisomatic boutons was quantified using the “Find spots close to surface” tool (ImarisXT extension) with a 0.8 µm distance threshold applied from the 3D-reconstructed NeuN+ surface.

For the *in situ* hybridization analysis, z-stacks (63X oil immersion objective, 1.4 NA, 2.2 digital zoom, 400 Hz, 1 µm step size) of 30-40 µm brain slices were imaged on an inverted Leica TCS-SP8 confocal (1024×1024 resolution, 16 bits) and analyzed using a custom macro in Fiji (ImageJ). Maximum z-stack projections were used for quantification. Somas of infected L5 IT and L5 ET neurons were drawn manually to create a mask of the soma surface. Cdh12 and Cdh13 RNA particles were detected automatically based on their intensity threshold, and a mask for each probe was generated using the “Analyze Particles” tool. *Cdh12* and *Cdh13* expression levels were determined as a percentage of the probe area normalized by the soma area.

For the monosynaptic tracing analysis, z-stacks (5 µm step size) were imaged on an inverted Zeiss ApoTome (10 X, 1280×800, 16 bits). For each mouse included in the analysis, 8 slices covering the entire S1 area were imaged, and infected cells were manually quantified using the Cell Counter tool in Fiji (ImageJ). The proportion of starter cells was determined as the number of receptor+/rabies+ cells normalized by the number of receptor+ cells across the 8 slices for both L5 pyramidal neuron subpopulations. The proportion of PV+ L5 IT and L5 ET inputs was determined as the number of PV+ inputs divided by the total number of inputs. The laminar proportion of PV+ inputs among L5 IT, L5 ET and L5 IT/ET inputs was determined as the number of PV+ inputs in each layer divided by the total number of PV+ inputs.

### In vitro patch-clamp recordings

Mice were deeply anesthetized with an overdose of sodium pentobarbital and transcardially perfused with 10 mL ice-cold slicing solution containing (in mM): 87 NaCl, 75 sucrose, 26 NaHCO_3_, 11 glucose, 7 MgCl_2_, 2.5 KCl, 1.25 NaH_2_PO_4_ and 0.5 CaCl_2_, oxygenated with 95% O_2_ and 5% CO_2_. The brain was quickly removed, and the injected hemisphere was glued to a cutting platform before being submerged in ice-cold slicing solution. 300 µm thick coronal slices containing S1Bf were cut using a vibratome (Leica VT1200S, Wetzlar, Germany) and incubated for 45-60 min at 32°C, and subsequently at room temperature, in the same solution. All salts were purchased from Sigma-Aldrich (St. Louis, MO). Slices were transferred to the recording setup and superfused with recording ACSF containing (in mM) 124 NaCl, 1.25 NaH_2_PO_4_, 3 KCl, 26 NaHCO_3_, 10 Glucose, 2 CaCl_2_, and 1 MgCl_2_, which was oxygenated with 95% O_2_ and 5% CO_2_ and heated to 34°C. Pipettes (3–5 MΩ) were made from borosilicate glass capillaries using a PC-10 pipette puller (P10, Narishige, London, UK). Miniature inhibitory postsynaptic currents (mIPSCs) were measured using an intracellular solution containing (in mM) 70 K-gluconate, 70 KCl, 10 Hepes, 4 Mg-ATP, 4 K2-phosphocreatine, and 0.4 GTP, adjusted with KOH to pH 7.3 (±290 mOsm). mIPSC recordings were performed at a holding voltage of -76 mV in the presence of 1 µM tetrodotoxin (TTX, HB1035) 10 µM 2,3-Dioxo-6-nitro-1,2,3,4-tetrahydrobenzo[f]quinoxaline-7-sulfonamide (NBQX, HB0443) and 50 µM D-(−)-2-Amino-5-phosphonopentanoic acid (D-APV, HB0225), all of which were purchased from Hello Bio (Bristol, UK). Recordings were made using a Multiclamp 700B amplifier (Molecular Devices, San Jose, CA). The signal was passed through a Hum Bug Noise Eliminator (Digitimer, Welwyn Garden City, UK), sampled at 10 kHz, and filtered at 3 kHz using a Digidata 1440A (Molecular devices, San Jose, CA). Cells were excluded if the access resistance (Ra) exceeded 30 MΩ. mIPSCs were analyzed using MiniAnalysis (SynaptoSoft, Decatur, GA, USA).

### Western blot

For Western blot analysis, HEK293T were rinsed with 1x ice-cold PBS. Samples were homogenized in lysis buffer containing 25 mM Tris-HCl pH 8, 50mM NaCl, 1% Triton X-100, 0.5% sodium deoxycholate, 0.001 % SDS and protease inhibitor cocktail (cOmplete Mini, Roche). Samples were resolved by SDS-PAGE and transferred onto PVDF membranes. Membranes were blocked with 5% Blotting-Grade Blocker (Bio-Rad, #1706404) in TBST (20mM Tris-HCl pH7.5, 150mM NaCl and 0.1% Tween20) for 1 hour. After incubation with chicken anti-HA (1:10 000, Abcam Ab1190) and mouse anti-actin (1:20 000, Sigma A3854) HRP-conjugated antibodies for 1h at room temperature, protein levels were visualized by chemiluminescence. Blots were scanned using a LI-COR Odyssey® Fc Imaging System.

### Statistical analysis

All statistical analyses were performed using GraphPad Prism 9 (GraphPad Software). Unless otherwise stated, parametric data were analyzed by *t*-test or one-way ANOVA followed by Holm-Sidak or Tukey *post hoc* analysis for comparisons of multiple samples. Non-parametric data were analyzed by the Mann-Whitney test, multiple t-test or Kruskal-Wallis one-way analysis followed by Holm-Sidak or Dunn *post hoc* analysis for comparisons of multiple samples. P values <0.05 were considered statistically significant. All data are presented as mean ± SEM.

